# Superpixel-ComBat multi-site harmonization of unpaired T1W MRI data in Huntington’s disease

**DOI:** 10.64898/2026.05.26.727928

**Authors:** Annabelle Coleman, Chang-Le Chen, Joelle Hanson-Baiden, Davneet S Minhas, Mahbaneh E Torbati, Charles M. Laymon, Sarah J Tabrizi, Edward J Wild, Dana L Tudorascu, Rachael I Scahill, Lauren M Byrne

**Author notes:** Joint senior authors (Rachael I Scahill, Lauren M Byrne). Correspondence to: Annabelle Coleman and Dr Lauren M Byrne, Full address: Huntington’s Disease Centre, UCL Queen Square Institute of Neurology, 2nd Floor Russell Square House, 10-12 Russell Square, London, WC1B 5EH.

## Abstract

**Background:** Pooling multi-site MRI data is essential for well-powered neuroimaging analyses, particularly in Huntington’s disease (HD), where large cohorts are needed to study disease-stage heterogeneity and subtle progressive neuroanatomical change. However, scanner-related variability hinders direct data pooling, confounding image-level methods such as voxel-based morphometry (VBM). Superpixel-ComBat (SP-ComBat), a voxel-level image-harmonization framework, effectively removes scanner effects but depends on traveling-subject data that are rarely available retrospectively. We extend SP-ComBat to unpaired multi-site T1-weighted MRI by introducing a pseudo-pairing framework that leverages demographically matched controls across scanners as surrogate traveling subjects.

**Methods:** Two pipelines were developed to estimate scanner effects under retrospective constraints: pipeline 1 used a small set of well-matched pseudo-pairs (n = 4) with bootstrap resampling to address scanners with limited sample sizes, while pipeline 2 used the recommended number of pseudo-pairs (n = 16) without resampling. Pseudo-pair images were parcellated into 3D-superpixels, and ComBat was applied within clusters to estimate scanner-specific adjustments for native-space harmonization. Pipeline performance was assessed in a representative multi-study dataset comprising six scanners from three HD cohorts (HD-YAS, HD-CSF, TRACK-HD; N = 144) and replicated in the full multi-study dataset (FMD; N = 548).

**Results:** Both pipelines improved image quality, aligned scanner-specific intensities, and preserved disease-related structural patterns. Pipeline 2 showed superior parameter re-estimation stability and was selected for the FMD. Harmonization eliminated systematic segmentation errors, enabled a single unified VBM pipeline across scanners, and increased sensitivity to HD-related voxel-wise atrophy.

**Conclusions:** SP-ComBat was effectively adapted for harmonization of unpaired multi-site structural MRI, reducing scanner bias while preserving biological variability and supporting unified VBM analyses across scanners.

## 1 Introduction

Large-scale neuroimaging studies and clinical trials increasingly depend on pooling structural magnetic resonance imaging (MRI) data across sites, yet inter-scanner variability - driven by differences in hardware design, magnetic field strength, manufacturer-specific implementations, and acquisition protocols (Preboske et al. 2005; Han et al. 2006; Takao et al. 2011; Takao et al. 2014; Raoul et al. 2025) - systematically affects image quality, producing variations in noise, contrast, and signal inhomogeneity (Chen et al. 2024). Consequently, scanner differences influence downstream analyses and often require site-specific pipeline optimization, hindering comparability and reproducibility across studies (Brown et al. 2020; Zhou et al. 2022; van Nederpelt et al. 2023; van Nederpelt et al. 2024). Without effective harmonization, such technical biases can obscure true biological signal and compromise the reliability of imaging-derived biomarkers (Hu et al. 2023).

One widely used technique vulnerable to these effects is voxel-based morphometry (VBM), a whole-brain imaging analysis method that identifies regionally specific structural brain differences (Ashburner and Friston 2000) without prior assumptions about the location, size, or shape of differences. VBM has been used extensively in neurological disorders, for example, in Huntington’s disease (HD) it has consistently demonstrated significant striatal grey-matter loss decades before clinical motor diagnosis (CMD) and progressive white-matter loss as the disease advances (Tabrizi et al. 2009; Minkova et al. 2018; Wang et al. 2024). However, because VBM operates on voxel-wise intensity patterns, even minor scanner-related signal inconsistencies can distort tissue segmentation and bias statistical inference (Hu et al. 2023). Conventional *post-hoc* covariate correction is often insufficient to account for these effects, motivating the development of dedicated harmonization strategies applied earlier in the neuroimaging pipeline.

Statistical harmonization is commonly implemented at the feature level. ComBat, an empirical Bayes method originally developed for genomics batch correction (Johnson et al. 2007), is widely applied to adjust derived measures from images such as cortical thickness (Fortin et al. 2018; Beer et al. 2020), regional volumes (Radua et al. 2020), or diffusion-tensor metrics (Fortin et al. 2017), to remove site effects while preserving biological signal. While multiple extensions of ComBat have been proposed (Orlhac et al. 2022; Hu et al. 2023), the influence of scanner differences varies across individual voxels within an image (Chen et al. 2020; Eshaghzadeh Torbati et al. 2021), meaning that a single global correction per feature cannot capture spatially heterogeneous scanner effects. This limitation has motivated image-level harmonization techniques that directly modify the voxel intensities (Radua et al. 2012; Radua et al. 2020; Zugman et al. 2022). The goal is to align structural MRI acquired from different scanners by reducing variations in image-related characteristics such as contrast- and signal-to-noise ratio (CNR, SNR, respectively), effectively rendering images comparable as if acquired under a single scanner (Chen et al. 2024; Yang et al. 2026). Examples include RAVEL, which removes unwanted between-scan variation at the voxel level (Fortin et al. 2016), and MISPEL, a supervised multi-scanner approach that also operates at the voxel level but requires paired data (Torbati et al. 2023). ComBat has also been adapted for image-level harmonization by applying the model to individual voxels in diffusion tensor imaging (Fortin et al. 2017). Although conceptually straightforward, this voxel-level approach is limited by the spatial consistency of voxels across subjects and generates a large number of estimated parameters, raising concerns about overfitting and statistical instability.

To overcome these limitations, Chen et al. (2024) introduced Superpixel-ComBat (SP-ComBat), which integrates ComBat modeling with 3D superpixel parcellation. Voxels are clustered by spatial proximity and intensity similarity using a simple linear iterative clustering algorithm (Achanta et al. 2012), and scanner-specific additive and multiplicative effects are estimated within each superpixel. The resulting parameter maps are projected back into native space to harmonize T1-weighted (T1W) MRI. Chen and colleagues (2024) found such harmonized images showed more consistent signal profiles and tissue contrasts across scanners, with substantial improvements in white matter and cerebrospinal fluid (CSF) SNR and increased consistency of grey matter SNR across sites. The coefficient of variation of grey matter volume was reduced by approximately 40%, demonstrating improved image quality and a marked reduction in scanner-related technical variability.

Despite these advantages, SP-ComBat and similar image-level harmonization methods rely on the availability of traveling subjects scanned across different sites to estimate scanner effects (Maikusa et al. 2021). The recommended 16–20 traveling subjects per scanner (Cetin Karayumak et al. 2019) are rarely available in retrospective studies, particularly in rare conditions such as HD. To address this gap, we developed a pseudo-pairing extension of SP-ComBat in which demographically-matched healthy controls from each scanner serve as surrogate traveling subjects. We applied this framework to unpaired structural T1W MRI from healthy controls and people with HD (pwHD) pooled from three retrospectively collected cohorts (HD Young Adult Study [HD-YAS], HD-CSF, TRACK-HD) spanning six scanners and implemented a unified VBM processing pipeline. Harmonization performance was evaluated using complementary qualitative and quantitative metrics encompassing image quality, residual scanner effects, and preservation of biological variability, as well as the impact on downstream morphometric analyses. This framework provides a foundation for robust whole-brain morphometric analyses in heterogeneous unpaired retrospective datasets and facilitates reliable investigation of neuroanatomical changes alongside biomarkers of interest.

## 2 Materials and methods

### 2.1 Study participants

We used baseline cross-sectional data from three retrospectively collected cohorts for harmonization (HD-YAS (Scahill et al. 2020), HD-CSF (Byrne et al. 2018), TRACK-HD (Tabrizi et al. 2009)), including healthy control participants, pwHD before CMD (pre-CMD) and after CMD (post-CMD). HD-YAS and HD-CSF were single-site studies, while TRACK-HD acquired imaging data across four international sites, together yielding six unique scanners for harmonization. Detailed protocols for each dataset have been previously published and are summarized in the Supplementary Methods 1.1 (Tabrizi et al. 2009; Byrne et al. 2018; Scahill et al. 2020). All participants provided written informed consent in accordance with the Declaration of Helsinki, and ethical approval was granted by local committees.

### 2.2 MRI acquisition and quality control

All T1W MRI data were acquired on 3-tesla (3T) scanners using 3D Magnetization Prepared Rapid Gradient Echo (MPRAGE) acquisition sequences (Supplementary Table 1). All scans underwent visual quality control (QC) before and after harmonization using MRView (https://www.mrtrix.org/), with raters blinded to scanner identity and disease status. Artifacts including motion, noise, truncation (also termed Gibbs ringing) and intensity inhomogeneity were rated using an original artifact rating scale (Supplementary Table 2), where higher scores indicated greater artifact burden; scans scoring above 3 were excluded.

### 2.3 SP-ComBat for unpaired inter-scanner variability estimation and harmonization

This cross-sectional study was pre-registered prior to analysis (10.17605/OSF.IO/S6TRK; Supplementary Figure 1). To model and remove inter-scanner variability at the image level, we adopted and extended the SP-ComBat framework (Chen et al. 2024) for retrospectively collected, unpaired multi-site data. Two alternative pipelines were developed, each estimating scanner effects using demographically matched healthy controls paired across scanners as surrogate traveling subjects (Figure 1). These pipelines were designed to address common constraints in retrospective studies, where it is often infeasible to simultaneously achieve strict demographic matching and the recommended sample size for image-level harmonization (∼16 pseudo-pairs per scanner; Cetin Karayumak et al. 2019).

**Figure 1:**
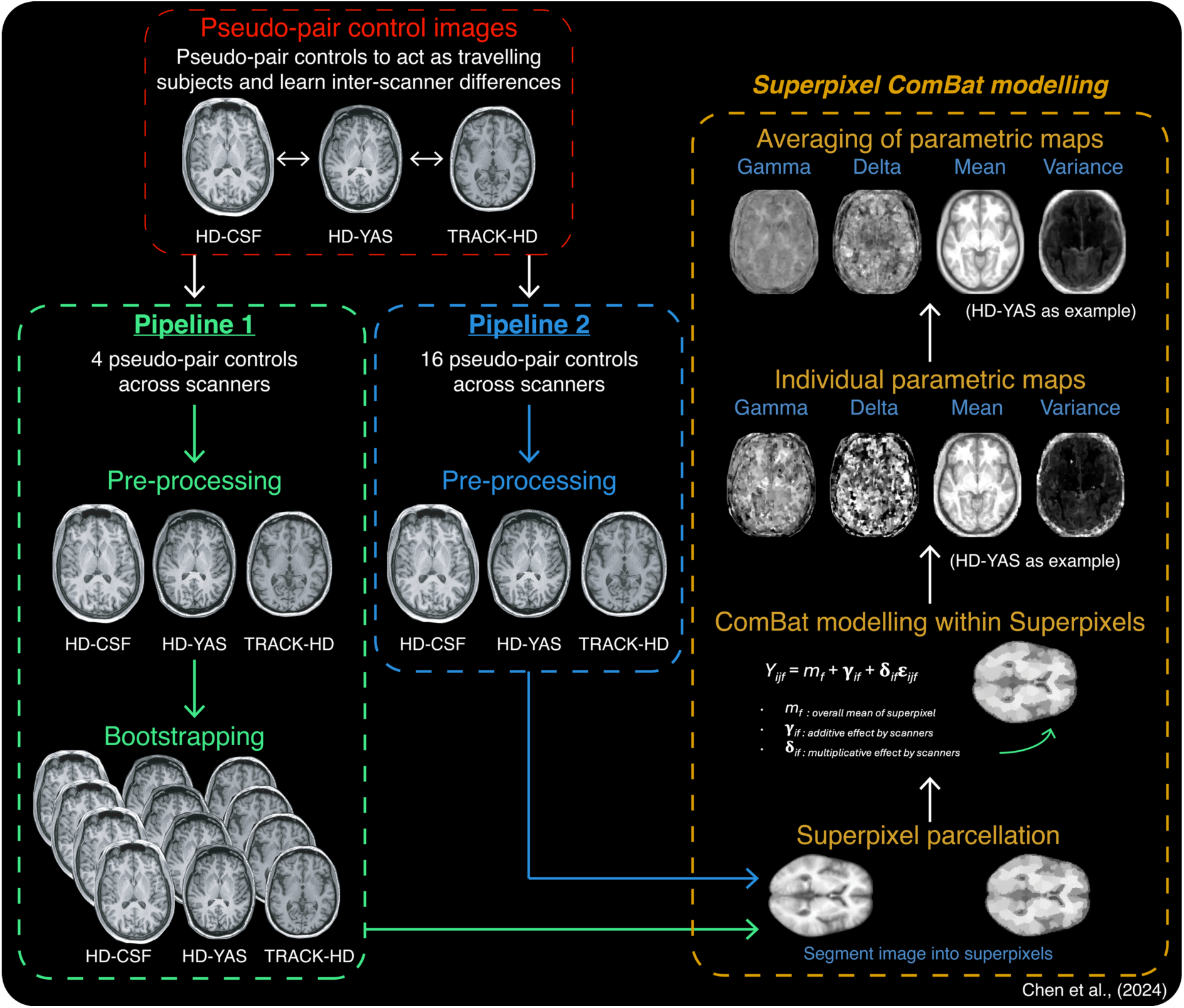
Overview of the unpaired SP-ComBat pseudo-pairing harmonization framework. Two pipelines were developed to estimate scanner effects using pseudo-paired healthy controls as surrogate traveling subjects. Pipeline 1 used four well-matched pseudo-pairs with an additional bootstrapping step to generate sufficient pseudo-pairs for scanner-effect estimation, whereas Pipeline 2 used 16 pseudo-pairs directly. Pseudo-pairs were pre-processed using CAT12 to standardize basic image quality and were registered to standard space through two-step spatial registration. SP-ComBat modelling followed Chen et al., 2024: 3D superpixel segmentation was applied to averaged pseudo-pair images, and ComBat modelling was used to estimate scanner effects within superpixels. Parametric maps were averaged at the group level and applied to perform site-effect removal for harmonization.

The first pipeline assessed harmonization feasibility under scanner sample-size constraints by selecting four well-matched pseudo-control pairs per scanner (Supplementary Table 3, Pipeline 1) and increasing the effective sample size via 25 iterations of bootstrap resampling to generate 25 synthetic pseudo-pairs for scanner-effect estimation. The second pipeline prioritized statistical power by matching the recommended number of pseudo-pairs per scanner (n = 16; Supplementary Table 3, Pipeline 2) without bootstrapping, accepting greater demographic variability. Each pipeline was applied first to a representative multi-study dataset (RMD), comprising balanced numbers of healthy controls, pwHD pre-CMD and post-CMD (RMD; n = 144; Supplementary Table 4), and harmonization performance was evaluated. The better-performing pipeline was subsequently used to harmonize the full multi-study dataset (FMD; n = 548; Supplementary Table 5), where evaluation was scaled-up in a larger cohort for further validation.

Across both pipelines, harmonization procedures included (Figure 1): (i) image preprocessing and spatial standardization, (ii) superpixel parcellation within pseudo-pairs for scanner-effect estimation with ComBat, and (iii) application of learned scanner differences to all images for native space scanner effect removal.

#### 2.3.1 Image pre-processing

Before harmonization, all T1-weighted MR images underwent standard preprocessing, as previously described in Chen et al., (2024) using SPM12 (v7771) and CAT12 (v12.9, Expert mode) packages implemented in MATLAB R2022b to standardize basic image quality and fulfill ComBat model assumptions. Briefly, all images were first resized using B-spline interpolation to ensure compatibility of image dimensions across scanners, followed by bias-field correction, denoising, and local intensity correction. Non-brain voxels were removed using a background mask to prevent formation of non-informative clusters during superpixel parcellation. Images were then subjected to global intensity normalization, rescaling voxel intensities to a range of 0–1000 based on the 99th percentile of the intensity distribution, facilitating comparison of image contrast across scanners while minimizing the influence of extreme intensity values. For inter-scanner variability estimation, images were spatially normalized using initial affine alignment and then nonlinear registration to MNI space using a diffeomorphic registration algorithm with geodesic shooting (Shoot registration; Ashburner and Friston 2011). This produced spatially normalized images for parameter estimation, together with forward and inverse deformation fields that were later applied to map SP-ComBat parameter estimates from standard space into each participant’s native image space.

#### 2.3.2 Superpixel-ComBat modelling

SP-ComBat modelling was used to estimate inter-scanner variability and perform site effect removal as previously described by Chen et al., 2024. Briefly, for SP-ComBat parameter estimation, spatially normalized pseudo-paired control images were averaged for superpixel parcellation. A three-dimensional superpixel segmentation was performed using a simple linear iterative clustering algorithm, which clusters voxels based on spatial proximity and intensity similarity (Achanta et al. 2012). Superpixel size constraints were applied to ensure each region contained sufficient voxels for stable statistical modelling (27 voxels minimum; Chen et al. 2024).

Within each superpixel, scanner-related effects were estimated using the ComBat empirical Bayes model (Johnson et al. 2007; Chen et al. 2024). Voxel intensities were modelled as the combination of a superpixel-specific mean intensity, scanner-specific additive (location; γ) effects, scanner-specific multiplicative (scale; δ) effects, and residual error. As scanner effects were learned using matched pseudo-paired control data, biological covariates were not included in the model. Parameter estimation was performed independently for each superpixel.

For each scanner, superpixel-wise additive and multiplicative parameter maps were generated and subsequently averaged across pseudo-paired samples to obtain group-level scanner-effect maps. These group-level maps were then used for harmonization. As parameter estimation was performed in MNI space, group-level maps were transformed into individual native space using participants’ inverse deformation fields derived during spatial registration. Scanner effects were removed at the image level by applying linear intensity correction using the estimated additive and multiplicative terms (Chen et al. 2024).

### 2.4 Voxel-based morphometry pre-processing

VBM analyses were performed using SPM12 to assess whole brain grey and white matter differences across unharmonized and harmonized T1W MRI. To enable valid comparison between unharmonized and harmonized images, both were processed using an identical pipeline, including the same templates, segmentation parameters, and spatial normalization procedures. This approach ensured that any observed voxel-wise differences reflected the effects of harmonization rather than preprocessing variability (Antonopoulos et al. 2023).

T1W images were segmented into tissue probability maps (TPMs) representing grey matter, white matter, CSF, bone, and soft tissue with preprocessing parameters: ‘medium (0.01)’ bias regularization, a 60-mm bias full-width-at-half-maximum (FWHM), two Gaussians per tissue class, and ‘thorough clean’ cleanup strength (Ashburner and Friston 2005). Grey and white matter TPMs were then nonlinearly aligned using Shoot registration which iteratively estimated a study-specific template based on the study population, enabling accurate inter-subject anatomical correspondence (Ashburner and Friston 2011). Following registration, TPMs were warped into MNI space and local tissue volumes were preserved with modulation (Ashburner and Friston 2000). Finally, the spatially normalized, modulated TPMs were smoothed with a 4-mm FWHM Gaussian kernel, consistent with the TRACK-HD study protocol (Tabrizi et al. 2009).

All pre-processing products were visually inspected blinded to scanner identity and disease status. Grey and white matter TPMs were visually examined for anatomical alignment to their corresponding T1W image. A segmentation error rating scale was used (Supplementary Table 6), with increasing numbers on the scale corresponding to decline in segmentation accuracy. TPMs that scored > 3 were excluded. If a TPM for one tissue class was excluded, all corresponding TPMs for that participant were excluded, both unharmonized and harmonized to ensure matched samples throughout.

Total intracranial volume (TIV) was delineated using semi-automated protocols on Medical Imaging Display and Analysis System (MIDAS; Freeborough et al. 1997), as previously described (Fox et al. 1996; Whitwell et al. 2001).

### 2.5 Statistical analysis

All analyses followed a pre-registered analysis plan (10.17605/OSF.IO/S6TRK) designed to assess whether SP-ComBat reduced scanner bias while preserving anatomical and disease-related variability. Statistical analyses were first performed on each harmonization pipeline in the RMD and compared to identify a suitable framework for unpaired SP-ComBat harmonization for the FMD. Criteria for pipeline selection are described in the Supplementary Methods 1.2 (Supplementary Table 7).

Unless otherwise stated, statistical analyses were performed using STATA version 19 (https://www.stata.com/) with a significance threshold of *p* < 0.05. Group differences in demographic variables were assessed within each scanner. Sex distributions were compared using chi-squared tests. CAG repeat length and disease burden score in HD groups, and age across all groups, were compared using one-way analysis of variance (ANOVA). Age was assessed between groups with post-estimation Wald tests.

#### 2.5.1 Image-quality assessments

Unharmonized and harmonized T1W images were evaluated using both qualitative and quantitative measures. Visual QC was performed side by side to identify any anatomical distortions introduced by SP-ComBat. Grey and white matter segmentations were overlaid on corresponding T1W images to examine segmentation consistency and assess whether harmonization affected tissue boundaries.

Quantitative image quality metrics on unharmonized and harmonized T1W MRI were derived using MRIQC (Esteban et al. 2017), including tissue-specific SNR, CNR, and coefficient of joint variation (CJV). SNR was estimated overall and separately for grey matter, white matter, and CSF, with higher values indicating better signal quality. CNR quantified grey-white matter contrast, with higher values reflecting improved tissue separability. CJV assessed head motion and intensity inhomogeneity artifacts, with higher values indicating poorer image quality. Each MRIQC metric was compared before and after harmonization using linear mixed-effects models including harmonization status, scanner, and their interaction as fixed effects, with subject-specific random intercepts. *Post-hoc* marginal effects were used to estimate overall harmonization effect and scanner-specific effects for each metric.

Changes in signal intensity distributions across scanners were examined visually using probability density plots of normalized intensities within intracranially masked regions to assess alignment of scanner-specific profiles while preserving tissue- and disease-related contrast.

#### 2.5.2 Voxel-wise morphometric analyses

VBM was conducted using SPM12 to assess the spatial extent of scanner- and disease-related effects of harmonization (Supplementary Methods 1.3; Supplementary Figure 2; Supplementary Table 8). General linear models with flexible factorial designs were specified, incorporating harmonization status and scanner or disease group as factors, as appropriate. To assess between disease-group differences, one-way ANOVA designs were specified separately in unharmonized and harmonized data. Age, sex, and TIV were included as covariates in all voxel-wise models. In unharmonized models, scanner was additionally included as a covariate. For each contrast, peak F/T-values, number of significant voxels/clusters, and spatial location of clusters were extracted and compared across harmonization status and pipelines.

#### 2.5.3 Volumetric analyses

Absolute grey and white matter volumes were extracted from modulated TPMs using the SPM “get_totals” utility, which computes tissue volume (mL) by summing voxel-wise probability values and scaling by voxel size. Volumes were analyzed using linear mixed-effects models with harmonization status, scanner, disease group, and their interactions as fixed effects, and subject-specific random intercepts to account for paired unharmonized and harmonized measurements per participant. Estimated marginal means and *post hoc* contrasts were used to quantify systematic absolute volume shifts attributable to harmonization across scanners and disease groups, as well as between-group differences in volumes normalized to TIV (volume/TIV × 100).

Agreement between harmonized and unharmonized absolute volumes was assessed using intraclass correlation coefficients (ICC), supplemented by Bland-Altman (Martin Bland and Altman 1986) and within-subject trajectory plots to visualize potential systematic bias introduced by harmonization.

#### 2.5.4 Additional assessments of harmonization effectiveness

Complementary analyses were conducted to further quantify residual scanner effects. First, principal component analysis (PCA) of intracranially masked voxel intensities was used to examine clustering by scanner before and after harmonization. Second, SP-ComBat parameters were re-estimated on harmonized images to assess whether scanner-specific additive and multiplicative effects approached theoretical null values. Third, convolutional neural network (CNN) classifiers were trained to predict scanner identity from unharmonized and harmonized images following the same protocol as Chen et al., (2024).

## 3 Results

### 3.1 Pseudo-paired controls without bootstrapping provided more stable scanner-effect estimation for unpaired SP-ComBat harmonization

We assessed two pipelines that used pseudo-paired healthy controls across scanners in the RMD to adapt SP-ComBat for unpaired data prior to application to the FMD (Figure 1). Across all assessments, both pipelines met QC criteria, comparably reduced scanner-related effects, and preserved biological variability (Supplementary Table 7). The pipelines diverged, however, on SP-ComBat re-estimation, where successful harmonization should drive re-estimated additive (γ) and multiplicative (δ) maps toward null values with minimal tissue-specific contrast.

Prior to harmonization, parameter maps reflected expected scanner-specific intensity signatures, with clear differences in tissue contrast and signal profiles across scanners in both pipelines (Figure 2A–D). Following harmonization, re-estimated parameters from Pipeline 1 exhibited only partial attenuation of additive effects and less stable multiplicative effects, particularly for HD-CSF (Figure 2E–F). In contrast, Pipeline 2 re-estimated additive-effect maps approached zero with minimal tissue-specific contrast, and multiplicative-effect maps were more spatially uniform across scanners (Figure 2G–H), indicating more complete removal of scanner-specific signal. Pipeline 2 was therefore selected for harmonization of the FMD on the basis of these findings and pre-defined selection criteria (Supplementary Table 7).

**Figure 2:**
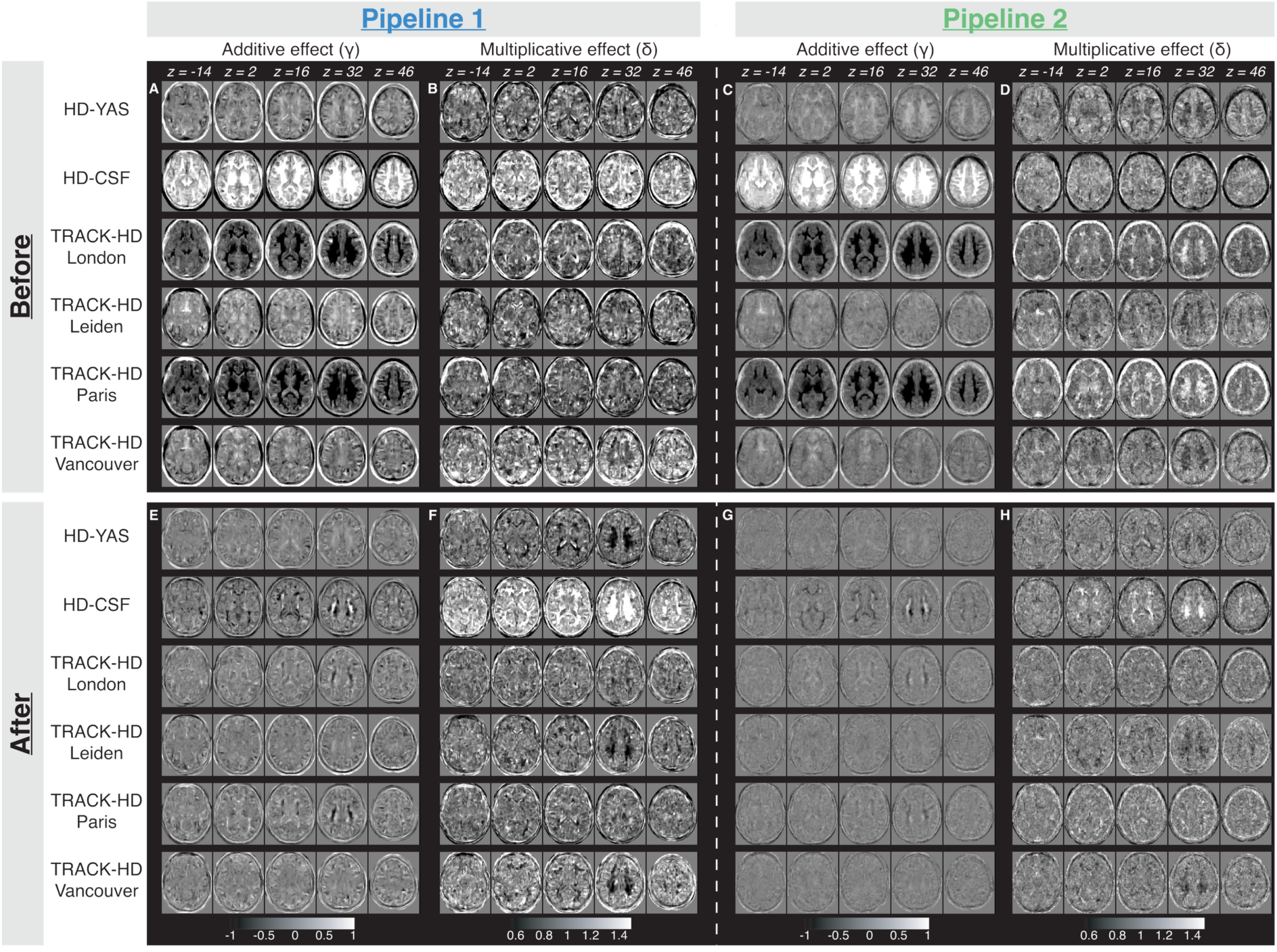
Scanner-specific additive and multiplicative parametric maps before and after harmonization for Pipeline 1 and Pipeline 2. Mean parametric maps derived from Pipeline 1 and Pipeline 2 showing additive (γ) and multiplicative (δ) effects at initial parameter estimation (A–D) and following pseudo-pair harmonization and re-estimation (E–H). Pipeline 1 maps were produced from all 25 bootstrapped pairs; Pipeline 2 maps were produced from all 16 pseudo-pairs. Maps are displayed in MRIcron with fixed intensity scales (γ: −1 to 1; δ: 0.5 to 1.5). z, axial slice position in MNI space. CSF, cerebrospinal fluid; HD, Huntington’s disease; HD-YAS, HD Young Adult Study; THD, TRACK-HD study site.

Raw scan QC for the RMD and FMD are detailed in Supplementary Results 2.1 and group demographics are displayed in Supplementary Tables 9-10. As harmonization assessment results were comparable across the RMD and FMD, subsequent results are presented for the FMD only.

### 3.2 Unpaired SP-ComBat improved image quality and removed inter-scanner variability

We applied Pipeline 2 to the FMD and firstly evaluated harmonization performance using qualitative visual inspection and quantitative MRIQC image quality metrics. Visual inspection confirmed that harmonization did not alter anatomical alignment or introduce spatial distortions, with post-harmonized scans displaying more consistent tissue contrast across scanners (Figure 3A). Quantitative metrics supported this. SNR increased significantly after harmonization for whole brain, grey matter, white matter, and CSF (all *p* < 0.0001; Figure 3B; Supplementary Table 11). CJV significantly decreased overall (adjusted mean difference [MD] = -0.08, 95% confidence interval [CI] -0.092 to -0.077, *p* < 0.0001), indicating reduced field inhomogeneity, although this reduction did not reach significance for the TRACK-HD Leiden scanner (Supplementary Table 11; MD = -0.016, *p* = 0.098). Global CNR did not change significantly (overall: MD = -0.018, CI -0.09 to 0.055, *p* = 0.623), though variance between scanners narrowed and significant scanner-specific gains emerged for HD-YAS, TRACK-HD Leiden, and TRACK-HD Vancouver (Supplementary Table 11).

**Figure 3:**
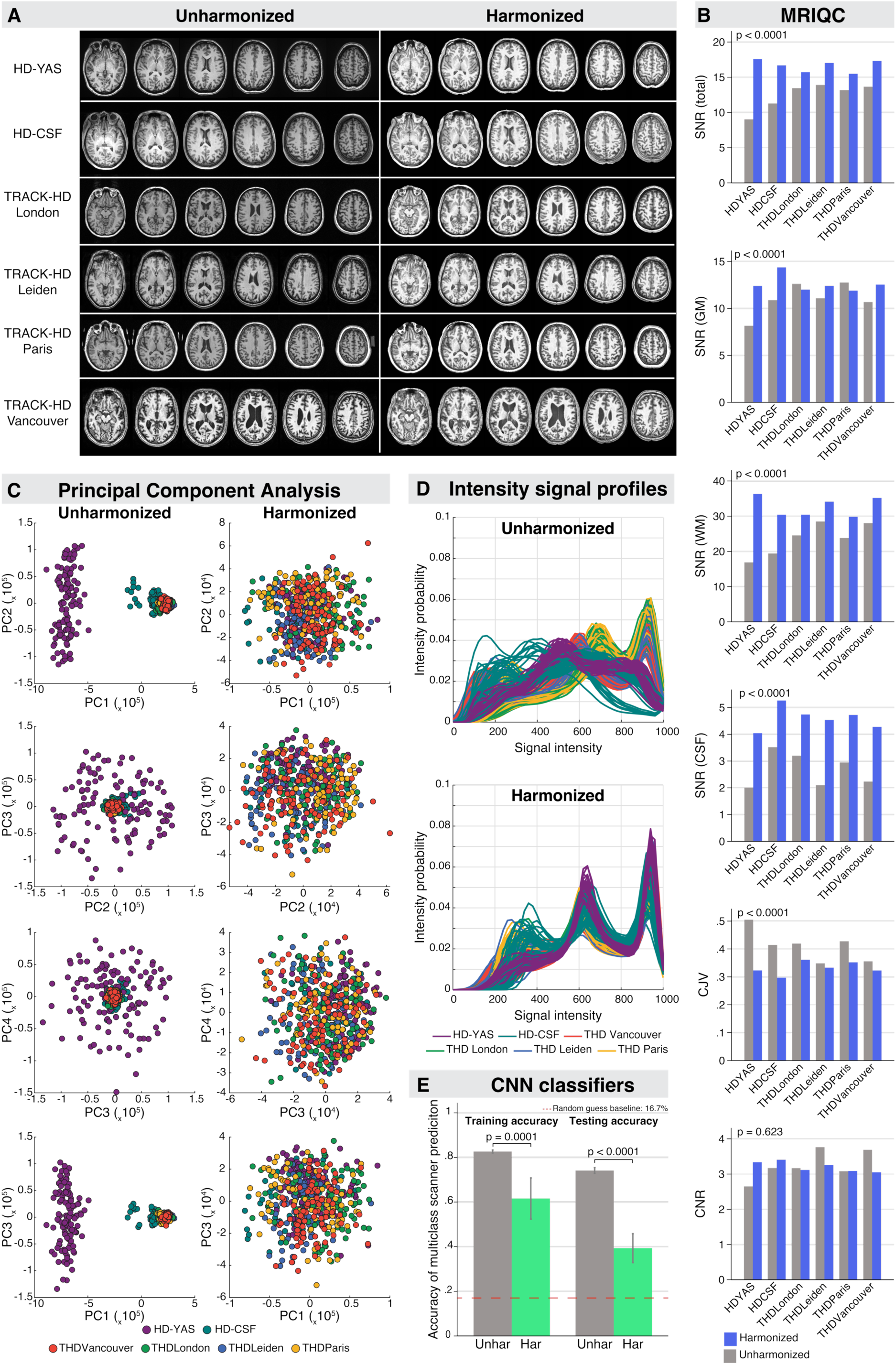
Image-level results before and after harmonization. A) Representative native-space T1W 3T MRI scans from each scanner before (unharmonized) and after (harmonized) SP-ComBat harmonization. B) Image quality metrics derived from MRIQC for unharmonized (grey) and harmonized (blue) images, including SNR for all tissue, grey matter, white matter, and CSF, and CJV and CNR; p-values reflect overall differences between conditions from linear mixed-effects models. C) PCA of intracranially masked voxels before (left) and after (right) harmonization. D) Probability density plots of normalized signal intensities within intracranially masked regions for unharmonized (top) and harmonized (bottom) images. E) Multiclass CNN classification accuracy for scanner prediction before and after harmonization, evaluated on training and test sets; the dashed line indicates the random-guess baseline for six scanners (16.7%); error bars represent 95% confidence intervals; p-values from two-sample t-tests. CJV, coefficient of joint variation; CNN, convolutional neural network; CNR, contrast-to-noise ratio; CSF, cerebrospinal fluid; GM, grey matter; har, harmonized; HD-CSF, HD Cerebrospinal Fluid Study; HD-YAS, HD Young Adult Study; PC, principal component; PCA, principal component analysis; SNR, signal-to-noise ratio; THD, TRACK-HD study site; unhar, unharmonized; WM, white matter.

To further characterise residual scanner effects beyond image quality metrics, voxel-intensity distributions were examined using PCA and intensity profiles, and scanner discriminability quantified using a multiclass CNN classifier (Figure 3C–E). Before harmonization, scanner identity dominated the variance structure, with PC1 explaining 97.61% of variance and HD-YAS and HD-CSF forming the most distinct clusters (Figure 3C; Supplementary Table 12), accompanied by highly variable voxel-wise signal intensity profiles across scanners (Figure 3D). After harmonization, scanner-specific clustering was markedly reduced, PC1 variance dropped to 35.68% (Figure 3C), and voxel-intensity profiles converged across scanners (Figure 3D). Repeating the PCA in healthy controls confirmed the reduction in PC1 variance reflected attenuation of scanner-related rather than disease-related signal (Supplementary Table 13; Supplementary Figure 3). CNN classification corroborated these findings: in the test set using images unseen during training, accuracy dropped from 0.741 to 0.393 following harmonization (*p* < 0.001), approaching the random-guess baseline for six scanners (16.7%). A corresponding reduction was observed in the training set (0.827 to 0.616; *p* = 0.001; Figure 3E). Together, these findings confirm effective attenuation of scanner-related signal while preserving raw image quality.

### 3.3 Unpaired SP-ComBat supported the use of a single VBM pipeline across diverse cohorts and scanners

We next examined whether this improvement in image-level consistency translated to more reliable application of a single VBM pipeline across all scanners. We did this by evaluating segmentation quality before and after harmonization using visual inspection and the segmentation error rating scale (Figure 4; Supplementary Table 6).

**Figure 4:**
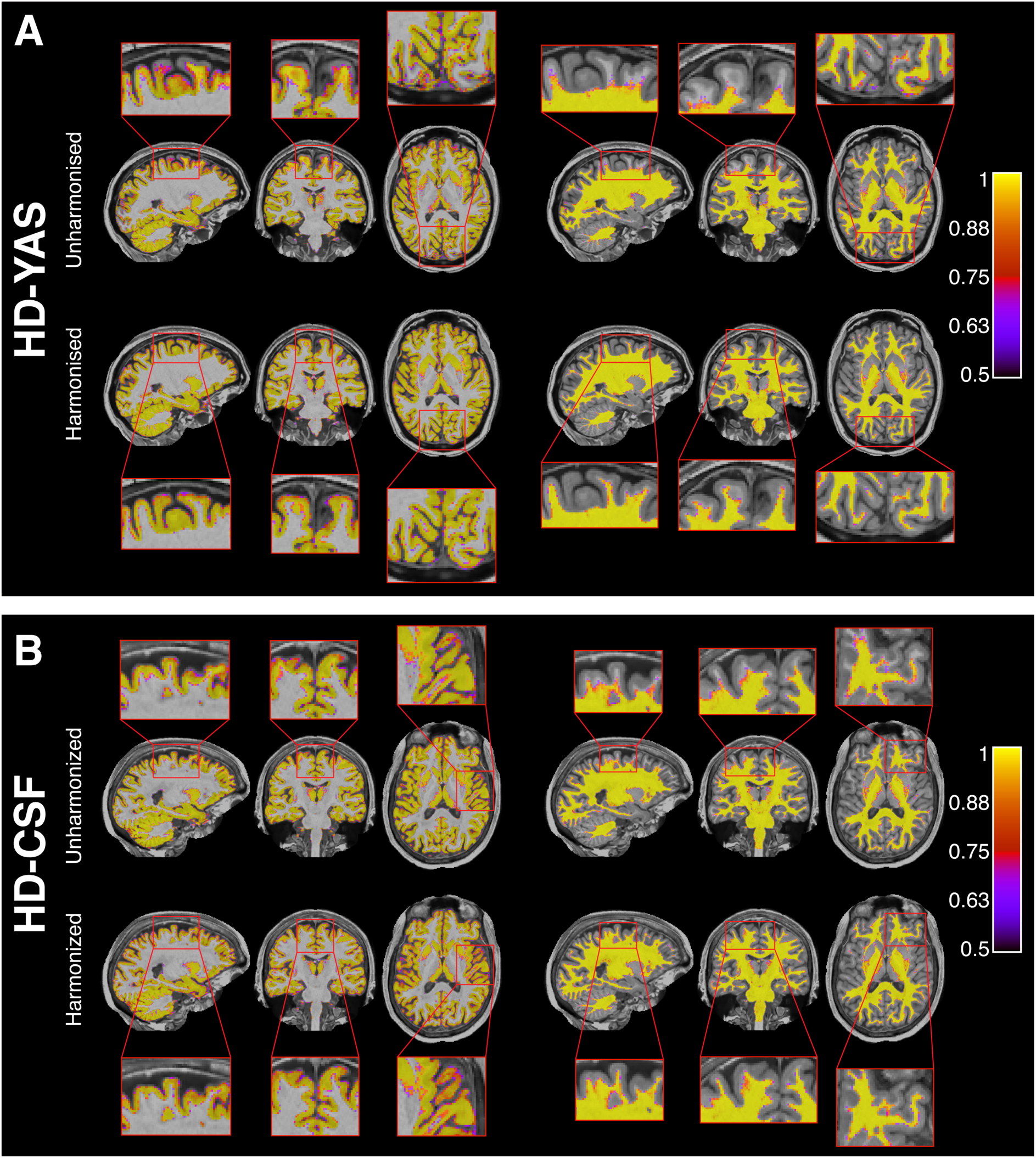
Effect of SP-ComBat harmonization on grey and white matter tissue segmentation in HD-YAS and HD-CSF scanners. Grey and white matter segmentations overlaid on native-space T1W MRI for representative HD-YAS (A) and HD-CSF (B) scans before and after harmonization. Images are displayed in three-plane orthogonal view with representative sagittal, coronal, and axial slices. Both unharmonized and harmonized segmentations are overlaid on the harmonized T1W MRI to enable direct visual comparison on a common reference image. HD, Huntington’s disease; HD-CSF, HD Cerebrospinal Fluid Study; HD-YAS, HD Young Adult Study.

When the unified SPM12 VBM pipeline was applied to unharmonized data, segmentation performance varied considerably by scanner (Supplementary Figures 4-5). Systematic errors were observed in HD-YAS and HD-CSF scanners, with HD-YAS the most severely affected: 97% of scans scored >3 on the segmentation error rating scale owing to systematic grey matter misclassification and upper cortical white matter under-segmentation (Figure 4A; Supplementary Table 14). HD-CSF showed a milder version of the same systematic pattern: 5% of scans failing for the same reasons (Figure 4B). For HD-CSF, a further 17% scored QC > 3 due to non-systematic errors, predominantly meningeal over-segmentation (Supplementary Table 14). By contrast, TRACK-HD scanners displayed no systematic segmentation bias but showed some non-systematic QC failures most commonly due to over-segmentation of the posterior meninges (Supplementary Table 14).

Following SP-ComBat harmonization, systematic errors were resolved in both affected HD-YAS and HD-CSF scans, driving systematic QC failure rates to zero (Supplementary Table 14; Figure 4). Segmentations showing non-systematic errors persisted after harmonization at comparable rates across all scanners, reflecting scanner-specific characteristics unrelated to harmonization (Supplementary Table 14).

To characterise the full extent of pipeline failure and its resolution with harmonization, unharmonized segmentations failing QC due to systematic errors were retained for the subsequent analyses; only segmentations failing due to non-systematic errors were excluded (Supplementary Table 14), leaving 501 matched unharmonized and harmonized TPMs taken forward for scanner-specific voxel-wise and volumetric analyses.

Voxel-wise comparisons between harmonized and unharmonized TPMs were performed to spatially quantify the extent of segmentation improvements and revealed significant cortical and subcortical clusters in which harmonized TPMs showed higher tissue probability values than unharmonized data (Supplementary Figure 6A–B; Supplementary Table 15). The largest differences were observed for HD-YAS, consistent with correction of the systematic segmentation errors in its unharmonized data (Supplementary Figure 6C; Supplementary Table 15; Figure 4A).

When assessing volumetric changes, harmonization produced systematic volumetric shifts consistent with correction of scanner-specific segmentation errors. Across most scanners, grey matter volume increased and white matter volume decreased (Grey matter: adjusted MD = 11.88 mL, 95% CI 9.29 to 14.47, *p* < 0.0001; white matter: adjusted MD = -17.62 mL, -19.25 to -15.98, *p* < 0.0001; Supplementary Figure 7; Supplementary Table 16), directionally consistent with correction of the grey matter misclassification and white matter under-segmentation seen in unharmonized HD-YAS scans. HD-CSF showed the opposite volumetric pattern (grey matter decreasing and white matter increasing), consistent with refinement of tissue boundaries in TPMs (Figure 4B). Voxel-wise spatial differences relative to HD-YAS were minimal (Figure 5D; Supplementary Figure 6), likely reflecting the milder and less spatially extensive nature of systematic errors in HD-CSF prior to harmonization. Despite these directional shifts, agreement between harmonized and unharmonized volumes remained excellent across all scanners with ICC exceeding 0.95 (Supplementary Figures 8-9 B-G) and Bland-Altman analyses stable across scanners (Supplementary Figures 8-9 H-M).

**Figure 5:**
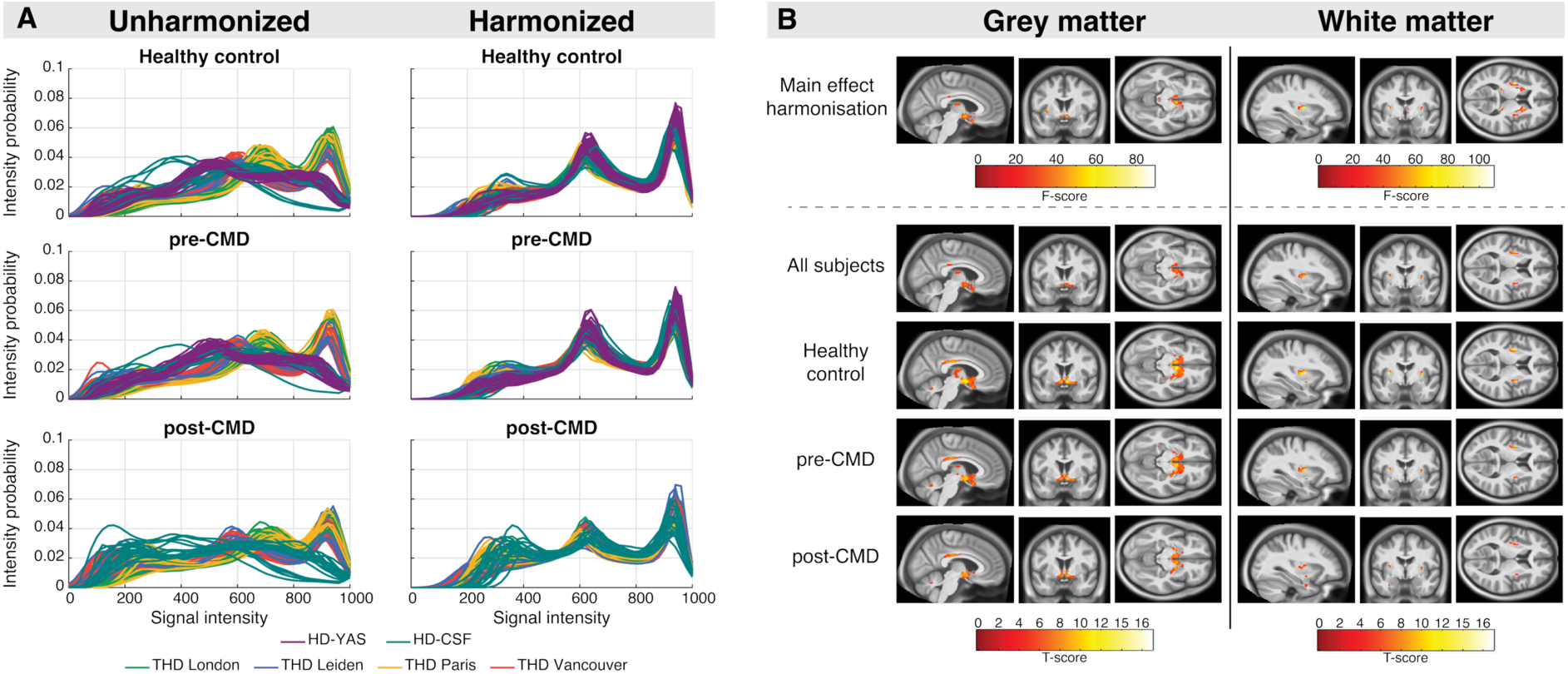
Signal intensity distributions and voxel-wise harmonization effects across disease groups. A) Probability density plots of normalized signal intensities within intracranially masked regions for unharmonized and harmonized scans, shown separately for healthy controls, pre-CMD, and post-CMD participants; colours denote individual scanners. B) Statistical parametric maps of grey and white matter differences between harmonized and unharmonized TPMs, shown for all participants passing QC and separately by disease group; significant clusters are overlaid onto sagittal, axial, and coronal sections of the study population-average brain; the main harmonization effect is displayed as F-scores and scanner-specific contrasts (harmonized > unharmonized) as T-scores; all comparisons are displayed on the same T-score scale (0–16); threshold p < 0.05, voxel-wise FWE-corrected (B). CMD, clinical motor diagnosis; FWE, family-wise error; HD, Huntington’s disease; HD-CSF, HD Cerebrospinal Fluid Study; HD-YAS, HD Young Adult Study; ICV, intracranial volume; post-CMD, pwHD after clinical motor diagnosis; pre-CMD, pwHD before clinical motor diagnosis; THD, TRACK-HD study site; TPMs, tissue probability maps.

### 3.4 Unpaired SP-ComBat preserved and enhanced sensitivity to HD-related neuroanatomical differences

We next assessed the effects of SP-ComBat harmonization on detection of biological variability. We began by examining whether harmonization effects were consistent across diagnostic groups, examining signal-intensity distributions, voxel-wise TPM patterns, and volumetric shifts separately for healthy controls, pwHD pre-CMD, and pwHD post-CMD. Voxel-wise and volumetric analyses were restricted to participants passing full segmentation QC, excluding both systematic and non-systematic QC failures, to ensure that observed effects reflected harmonization rather than residual segmentation error (final N = 379; Supplementary Table 14).

Harmonization aligned signal-intensity distributions across scanners comparably in healthy controls, pre-CMD, and post-CMD (Figure 5A); greater variability at low intensities in pwHD post-CMD was expected given the ventricular enlargement characteristic of advanced disease and was present both before and after harmonization, confirming it reflects underlying biology rather than a harmonization artifact. Direct voxel-wise comparisons of harmonized and unharmonized TPMs showed consistent spatial patterns of increased tissue probability values across all disease groups (Figure 5B; Supplementary Table 17). Volumetric shifts were similarly modest and directionally uniform across groups: grey-matter volumes increased (adjusted MD = 4.2-8.4 mL) and white-matter volumes decreased (adjusted MD = -12.6 to -6.8 mL; Supplementary Figure 10; Supplementary Table 18), with agreement between harmonized and unharmonized volumes high across groups (ICC > 0.9; Supplementary Figure 10).

We then assessed whether harmonization preserved and enhanced sensitivity to HD-related neuroanatomical differences. Voxel-wise disease-group comparisons revealed known patterns of HD-related grey and white matter atrophy were preserved after harmonization and, in several regions, showed greater spatial extent and higher peak T-values than unharmonized data (Figure 6A,C; Supplementary Table 19). This was most evident in the comparison between pwHD pre-CMD and healthy controls, where bilateral striatal atrophy, a canonical early marker of HD neurodegeneration, showed a marked increase in spatial extent and magnitude after harmonization, most notably in the right putamen (unharmonized: 1,489 voxels, peak T = 11.67; harmonized: 3,862 voxels, peak T = 14.51; Figure 6A,C; Supplementary Table 19). In the comparison between pwHD post-CMD and healthy controls, widespread grey and white matter atrophy was observed both before and after harmonization (Figure 6A,C; Supplementary Table 19-20). Volumetric analyses showed that disease-stage differences were statistically consistent before and after harmonization across all comparisons, with harmonization yielding slightly more conservative estimates of mean volumetric differences (Figure 6B,D, Supplementary Table 19).

**Figure 6:**
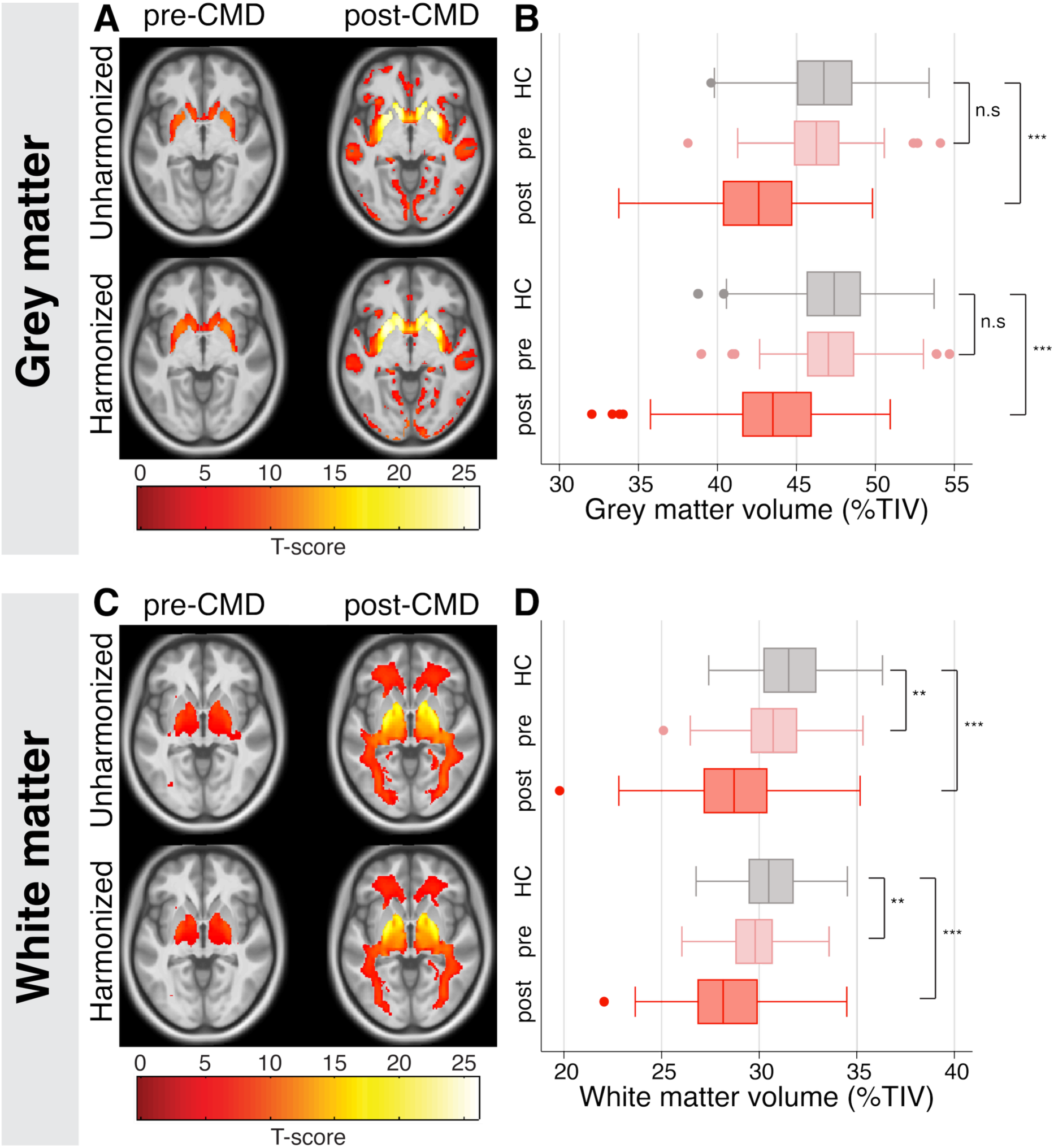
Voxel-wise and volumetric disease group differences before and after SP-ComBat harmonization. Statistical parametric maps of grey matter (A) and white matter (C) differences between disease groups in unharmonized and harmonized TPMs, restricted to participants passing segmentation QC. T-contrasts compare grey and white matter values in pwHD pre-CMD and post-CMD relative to healthy controls; significant clusters are overlaid onto axial sections of the study population-average brain; threshold p < 0.05, voxel-wise FWE-corrected; all maps are displayed on the same T-score scale (0–25). Volumetric differences in grey (B) and white matter (D) before and after harmonization, stratified by disease group; volumes are adjusted for TIV (GM/TIV × 100); boxes show the first and third quartiles, the central band shows the median, and whiskers extend to 1.5 times the interquartile range from the first and third quartiles; p-values are from Bonferroni-corrected post-hoc pairwise comparisons of estimated marginal means from a mixed-effects model (fixed effects: harmonization × scanner × disease group; random intercept: subject). **p < 0.01, ***p < 0.0001. CMD, clinical motor diagnosis; FWE, family-wise error; HC, healthy control; HD, Huntington’s disease; n.s., not significant; post-CMD, pwHD after clinical motor diagnosis; pre-CMD, pwHD before clinical motor diagnosis; TIV, total intracranial volume; TPMs, tissue probability maps.

## 4 Discussion

Leveraging data from 538 participants across six scanners, we show that image-level harmonization with SP-ComBat can be extended to unpaired, retrospectively collected multi-site T1W MRI, mitigating scanner-related variability while preserving anatomical integrity and biologically meaningful disease effects. Unlike feature-level harmonization approaches applied to derived morphological metrics (Fortin et al. 2018; Hu et al. 2023; Yang et al. 2026), image-level harmonization directly stabilizes intensity distributions prior to downstream analyses, thereby mitigating scanner bias throughout the analytical pipeline (Chen et al. 2024). Consequently, SP-ComBat improved image quality, resolved systematic tissue misclassification, and enhanced voxel-wise sensitivity to canonical patterns of HD neurodegeneration within a single unified framework.

A persistent challenge in multi-site neuroimaging is the dependence on traveling-subject data to estimate scanner effects at the image level (Hu et al. 2023). This limitation is especially problematic in rare diseases such as HD, where data are typically acquired in site-specific studies designed around individual research questions, making retrospective pooling across cohorts inherently difficult. Our pseudo-pairing extension of SP-ComBat removes this requirement by using demographically matched healthy controls as surrogate traveling subjects, enabling image-level harmonization of retrospectively collected multi-site data. Comparing two pseudo-pairing strategies clarified the trade-off between matching precision and sample size. Bootstrapping a small, well-matched set (Pipeline 1, four pairs per scanner) inflated uncertainty in additive and multiplicative parameter estimates, whereas using the recommended sample directly (Pipeline 2; n = 16; Cetin Karayumak et al. 2019) yielded more stable estimates and more complete attenuation of scanner-related signal, even at the cost of looser demographic matching. This suggests that the bootstrapping procedure introduced variability into scanner-effect estimation, likely reflecting the stochastic nature of resampling, where occasional biased draws can affect parameter stability. Pipeline 2’s direct use of the recommended sample size avoided this dependence and yielded more reliable estimates, although future work characterising individual bootstrap iterations could clarify when resampling-based approaches remain useful.

Applying pipeline 2 to the full dataset, this approach attenuated scanner-driven signal while leaving biologically meaningful variability intact across multiple converging analyses. Improvements in tissue-specific SNR and reduced CJV indicated lower noise and field inhomogeneity, while preserved global CNR and narrowed between-scanner variance showed that tissue contrast itself was not flattened. The same pattern emerged from data-driven analyses: scanner identity, which dominated voxel-intensity variance before harmonization (PC1 = 97.6%), no longer organized the data afterwards (PC1 = 35.7%), and a CNN trained to discriminate scanners performed closer to chance on harmonized images. Together, these analyses indicate that the corrections selectively removed scanner-driven signal. Notably, the corrections themselves were not uniform, reflected by distinct volumetric shifts across scanners following harmonization. This adaptive behavior is consistent with the empirical Bayes framework underlying ComBat, which estimates scanner-specific additive and multiplicative effects to align data toward shared statistical properties rather than imposing a global correction (Johnson et al. 2007; Fortin et al. 2017; Chen et al. 2024). Each scanner’s data is therefore adjusted according to its own estimated bias, avoiding the overcorrection that a uniform adjustment would impose on scanners with distinct intensity profiles.

The benefits of these intensity-level corrections translated directly into downstream morphometric accuracy, fundamentally changing how the data could be processed. With unharmonized data, applying a single processing pipeline across scanners was not feasible. While SPM12 accurately segmented tissue classes across most scanners, it systematically failed on the HD-YAS scans. CAT12, another VBM software, can segment these scans successfully (Scahill et al. 2020), but adopting different software for one site would have required mixing methodologies across the wider study and would have introduced pipeline-dependent variability in results (Antonopoulos et al. 2023). This exposed a known central challenge in multi-site imaging studies: without image-level harmonization, researchers are often required to tailor pipelines to individual scanners, which in turn compromises the interpretability and comparability across sites (Zhou et al. 2022). By stabilizing image intensities prior to processing with SP-ComBat, these segmentation failures were eliminated and site-specific optimization became unnecessary, allowing a single unified VBM pipeline to be applied across all datasets. The scale of this recovery was substantial: applying a single unified pipeline to unharmonized data would have yielded only 384 grey matter and 414 white matter TPMs passing strict QC (score < 3), whereas harmonization increased these to 505 and 536 respectively, recovering approximately 30% of otherwise-excluded scans. This enabled heterogeneous multi-site data to be treated as a single large cohort, increasing statistical power while eliminating pipeline-induced methodological bias.

Beyond the methodological gains, a key concern in any harmonization framework is whether the correction itself introduces disease-specific bias or dampens biologically meaningful signal (Hu et al. 2023). In this study, harmonization effects were consistent across disease groups: intensity, voxel-wise, and volumetric shifts induced by SP-ComBat did not vary by disease status, suggesting no differential bias related to advanced HD phenotype. Importantly, biological signal was preserved, as voxel-wise and volumetric group differences between pwHD and controls remained evident after harmonization. In fact, harmonization increased sensitivity to HD-related neurodegeneration in voxel-wise analyses, reflected by higher peak T-values across all comparisons. This was most evident in pwHD pre-CMD, where a broader spatial extent of bilateral striatal atrophy emerged despite no significant differences in absolute grey matter volumes.

This dissociation between voxel-wise and volumetric findings may reflect how harmonization-driven SNR improvements affect the two scales of measurement differently. In unharmonized data, tissue segmentation was less precise at boundaries, where voxel tissue class classification was more uncertain. These small misclassifications accumulate, inflating variability and leading to over- or under-estimation of tissue volumes. Following harmonization, improved SNR enhanced the contrast between tissue classes, allowing the segmentation algorithm to assign voxels with greater confidence to either grey or white matter. This resulted in cleaner boundaries and reduced the inclusion of spurious, uncertain voxels. Consequently, volumetric estimates derived from harmonized data were tighter and more conservative. At the voxel level, by contrast, the same reduction in scanner-related noise sharpened detection of focal biological effects rather than averaging them away (Hu et al. 2023; Chen et al. 2024). Together, these findings suggest that voxel-wise sensitivity gains from harmonization within larger multi-study datasets may enhance detection of early unbiased neurodegenerative changes in pwHD (Scahill et al. 2020; Tabrizi et al. 2022).

Realising this clinical potential will depend on addressing several limitations. The pseudo-pairing approach depends on the availability of a sufficient number of well-matched controls per scanner to ensure stable estimation of scanner-specific effects. This may limit harmonization in scanners with small sample sizes; therefore, alternative data-augmentation strategies or alternative pairing approaches may be required and warrant systematic evaluation. A related concern is the absence of traveling-subject data: while multiple complementary analyses demonstrated attenuation of scanner effects, the absence of true traveling-subject data precludes direct validation against a formal gold standard. Future work applying the pseudo-paired framework to datasets with traveling subjects will be important for further validation. The present study used a cross-sectional harmonization pipeline; extension to longitudinal designs will be necessary for modelling within-subject disease trajectories, which requires accounting for potential interactions between scanner effects and time. Finally, the current implementation was evaluated using research-grade 3T T1W MRI; defining its performance across other field strengths, additional modalities, and more heterogeneous acquisition protocols will be required for establishing the boundaries of generalizability in large-scale neuroimaging consortia (Chen et al. 2024).

In conclusion, unpaired SP-ComBat provides a scalable framework for image-level harmonization of multi-site neuroimaging data, particularly suited to rare diseases where prospective harmonization and traveling-subject acquisitions are not feasible. By reducing scanner-related bias while preserving anatomical and biological validity, this approach enabled unified analysis of multi-scanner datasets and enhanced sensitivity to disease-related neurodegeneration.

## 5 Data availability

The data analyzed during this study are available from the corresponding author upon reasonable request.

## 7 CRediT authorship contribution statement

Annabelle Coleman: conceptualization, data curation, formal analysis, investigation, methodology, validation, visualization, writing original draft, writing review & editing

Chang-Le Chen: conceptualization, methodology, software, validation, writing review & editing

Joelle Hanson-Baiden: methodology, validation, writing review & editing

Davneet S Minhas: software, validation, writing review & editing

Mahbaneh E Torbati: software, writing review & editing

Charles M. Laymon: software, writing review & editing

Sarah J Tabrizi: resources, supervision, writing review & editing

Edward J Wild: resources, supervision, writing review & editing

Dana L Tudorascu: software, supervision, writing review & editing

Rachael I Scahill: methodology, resources, supervision, visualization, writing review & editing

Lauren M Byrne: conceptualization, funding acquisition, methodology, resources, supervision, visualization, writing original draft, writing review & editing

## 9 Acknowledgements

The authors gratefully acknowledge all participants in the TRACK-HD, HD-YAS, and HD-CSF studies for their invaluable contributions to this research.

