## Supplementary Tables and Figures for "Superpixel-ComBat multi-site harmonization of unpaired T1W MRI data in Huntington’s disease"

#### Supplementary materials

Annabelle Coleman[1], Chang-Le Chen[2], Joelle Hanson-Baiden[1], Davneet S Minhas[3], Mahbaneh E Torbati[4], Charles M. Laymon[2,3], Sarah J Tabrizi[1,5], Edward J Wild[1], Dana L Tudorascu[6,7], Rachael I Scahill[1†], Lauren M Byrne[1†]

Author affiliations:

1 Huntington's Disease Centre, Department of Neurodegenerative disease, UCL Queen Square Institute of Neurology, University College London, London, UK

2 Department of Bioengineering, Swanson School of Engineering, University of Pittsburgh, Pittsburgh, PA, USA

3 Department of Radiology, University of Pittsburgh School of Medicine, Pittsburgh, PA, USA

4 Department of Anesthesiology, Perioperative and Pain Medicine, Stanford University, Stanford, CA, USA

5 UK Dementia Research Institute, University College London, London, UK

6 Department of Psychiatry, School of Medicine, University of Pittsburgh, Pittsburgh, PA, USA

7 Department of Biostatistics and Health Data Science, University of Pittsburgh, Pittsburgh, PA, USA

[†] Joint senior authors

Correspondence to: Annabelle Coleman and Dr Lauren M Byrne

Full address: Huntington's Disease Centre, UCL Queen Square Institute of Neurology, 2nd Floor Russell Square House, 10-12 Russell Square, London, WC1B 5EH

### 1 Supplementary methods

#### 1.1 Participant cohorts

Data from three retrospectively collected cohorts were used for harmonization (HD-YAS, HD-CSF, TRACK-HD). HD-YAS is an ongoing single-site longitudinal study investigating the earliest manifestations of HD pathology in pwHD ~24 years from predicted CMD at baseline (Scahill et al. 2020; Scahill et al. 2025). In 2018, 67 healthy controls and 64 pwHD underwent clinical and cognitive testing, 3-tesla (3T) volumetric MRI, and biofluid collections (blood and CSF). pwHD required a positive HD genetic test with CAG ≥40, disease burden score (DBS: age x (CAG-35.5); Penney et al. 1997) < 240, and Unified HD Rating Scale (UHDRS) total motor score (TMS; Kieburtz 1996) < 5. Healthy controls were age- and sex-matched gene-negative individuals.

HD-CSF was a single-site two-year longitudinal investigation of CSF in 80 participants (20 healthy controls, 20 pre-CMD, and 40 post-CMD) who had matched 3T T1W MRI and blood collection (Byrne et al. 2018; Rodrigues et al. 2020). Pre-CMD participants had CAG ≥40 with diagnostic confidence score (DCS; Kieburtz 1996) of < 4. Early post-CMD participants had CAG ≥40, DCS = 4, and were staged by UHDRS total functional capacity (TFC; Martinez-Martin et al. 2017) score (Stage 1, TFC 11-13; Stage 2, TFC 10-7; Stage 3, TFC 7-10).

TRACK-HD was a four-year multi-site observational study evaluating biomarkers of disease progression in 366 participants (healthy controls, pre-CMD, and post-CMD) across four sites: London; Leiden; Paris; and Vancouver (Tabrizi et al. 2009; Tabrizi et al. 2011; Tabrizi et al. 2012; Tabrizi et al. 2013). Annual assessments included 3T T1W MRI, blood sampling, and clinical, motor, and cognitive assessments. All pwHD had a CAG ≥40. Pre-CMD participants required DBS >250 and a TMS ≤5, indicating no substantial clinical motor signs. Post-CMD participants had a DCS of 4, and were within Shoulson and Fahn stage 1 or 2 (TFC ≥7; Shoulson and Fahn 1979). Healthy controls were age- and sex-matched spouses, partners, or gene negative siblings.

Detailed acquisition protocols for each dataset have been previously published (Tabrizi et al. 2009; Byrne et al. 2018; Scahill et al. 2020).

#### 1.2 Criteria for pipeline selection

The accumulation of results from each pipeline were evaluated and a decision was made about which pipeline to choose for larger-scale analysis (Supplementary Table 7). The aim was to identify a pipeline that most effectively mitigated scanner-related bias while preserving biologically meaningful variability and anatomical integrity. Each pipeline was evaluated across four key domains: (1) scanner bias reduction; (2) biological variability preservation; (3) structural integrity; (4) computational feasibility. While the ideal design would excel across all evaluation domains, trade-offs may be necessary. In cases where designs perform similarly, preference was given to feasible or more computationally efficient options. If none of the evaluated experimental designs met an acceptable standard due to persistent distortions or inadequate scanner effect removal, a standard unharmonized pipeline would be used with scanner as a covariate (no harmonization procedures).

#### 1.3 VBM statistical analysis contrast design

Voxel-wise comparisons on both grey and white matter between unharmonized and harmonized TPMs were conducted to identify any systematic changes in brain morphology introduced by SP-ComBat. We anticipated this whole-brain unbiased voxel-wise approach would provide information on regional biases; uncover specific brain regions that were more impacted by harmonisation due to scanner effects or disease-specific morphological differences. All voxel-wise analyses were conducted using SPM12.

##### Flexible factorial design specification

For within scanner and within disease comparisons, a general linear model (GLM) with a flexible factorial design questioned if harmonization impacted the resulting grey and white matter values, and whether any influence was dependent on a specific scanner of disease group. Two separate GLMs were constructed: one for grey matter TPMs and one for white matter TPMs. The design matrix included two predictors (Supplementary Figure 2A-B): (1) harmonization (two levels: unharmonized and harmonized) and (2) scanner (six levels: HD-YAS, HD-CSF, TRACK-HD London, TRACK-HD Leiden, TRACK-HD Paris, and TRACK-HD Vancouver) or disease group (three levels: controls, pre-CMD, and post-CMD). Covariates included age, sex, and TIV.

##### ANOVA design specification

To assess disease group differences in the unharmonized and harmonized datasets, independent voxel-wise one-way ANOVAs were built to question if there were significant differences between grey/white matter regions between each disease group. Four different one-way ANOVAs were performed: unharmonised grey (1) and white (2) matter TPMs, harmonized grey (3) and white (4) matter TPMs. The design matrix included one predictor (Supplementary Figure 2C): disease group (three levels: controls, pre-CMD, and post-CMD). Covariates included age, sex, and TIV. Scanner was added as a covariate in the unharmonized ANOVA.

##### Contrast specification

GLMs were estimated using the 'Estimate' batch job in SPM12. To examine scanner- and disease-specific harmonisation effects or disease group differences, the ‘Results’ batch job was used to define F- and T-contrasts (Supplementary Table 8). For the effects of harmonization, F-contrasts were specified to identify regions where harmonisation significantly altered grey/white matter values across all scanners or disease groups, while T-contrasts were used to detect scanner- or disease- specific increases or decreases in grey/white matter values due to harmonisation. For disease group differences comparing pwHD to healthy controls, T-contrasts were specified to identify regions with significantly different grey/white matter values between healthy controls and pre-CMD participants, and healthy controls and post-CMD participants.

### 2 Supplementary results

#### 2.1 T1W MRI QC

All pseudo-paired control T1W MRI scored 0 on the QC severity scale with no artifacts detected.

For the RMD, 127/144 (88%) scans scored 0 on the severity scale with no artifacts detected, 12/144 (8%) scored 1 with minor artefacts with half being from HD-CSF scanner, and 5/144 (5%) scored 2 with mild artefacts that were slightly more prominent. 4/5 of the scans that were rated a score of 2 were from the post-CMD HD group. Although some artifacts persisted, none were deemed bad enough for exclusion. After harmonization, for both pipelines the visual QC outcomes were identical to those of the unharmonized T1W MRI.

For the FMD, 469/548 (86%) scans scored 0 on the QC severity scale with no artifacts detected, 41/548 (7%) scored 1 with minor artefacts, 46/548 (8%) scored 2 with mild artefacts that were slightly more prominent, 5/548 (1%) scored 3 with moderate artefacts, and 7/548 (1%) scored 4 and above with heavy artefacts detected (excluded). All scans that were rated 3 or higher were pwHD post-CMD. Three scans were excluded due to the presence of non-HD pathology. This gave an exclusion rate of 1.8% in the raw T1W QC, leaving 538 scans for harmonization. After harmonization, the same QC results were replicated as before harmonization with all $n$ = 538 passing visual QC.

### 3 Supplementary Tables

*Supplementary Table 1: MRI acquisition parameters for each scanner across the three study cohorts*

| Study | HD-YAS | HD-CSF | TRACK-HD London | TRACK-HD Leiden | TRACK-HD Paris | TRACK-HD Vancouver |
|---|---|---|---|---|---|---|
| **Location** | London | London | London | Leiden | Paris | Vancouver |
| **Scanner** | Siemens Prisma | Siemens Prisma | Siemens Trio | Philips Achieva | Siemens Trio | Philips Achieva |
| **Field (T)** | 3 | 3 | 3 | 3 | 3 | 3 |
| **Sequence** | T1W 3D MPRAGE | T1W 3D MPRAGE | T1W 3D MPRAGE | T1W 3D MPRAGE | T1W 3D MPRAGE | T1W 3D MPRAGE |
| **Plane** | Sagittal | Coronal | Sagittal | Sagittal | Sagittal | Sagittal |
| **Voxel size ($mm^3$)** | 1x1x1 | 1x1x1 | 1x1x1 | 1x1x1 | 1x1x1 | 1x1x1 |
| **ST (mm)** | 1 | 1 | 1 | 1 | 1 | 1 |
| **TR (ms)** | 2530 | 2000 | 2200 | 7.7 | 2200 | 7.7 |
| **TE (ms)** | 3.34 | 2.05 | 2.2 | 3.5 | 2.2 | 3.5 |
| **TI (ms)** | 1100 | 850 | 900 | 950 | 900 | 950 |
| **FA (°)** | 7 | 8 | 10 | 8 | 10 | 8 |
| **FOV ($mm^3$)** | 256 | 240 | 280 | 240 | 280 | 240 |
| **Matrix size** | 224x224 | 256x240 | 256x256 | 224x224 | 256x256 | 224x224 |

Scanner specific acquisition details outlined in retrospective studies (Tabrizi et al. 2009; Byrne et al. 2018; Scahill et al. 2020).
FA, flip angle; FOV, field of view; HD, Huntington's disease; HD-CSF; HD Cerebrospinal Fluid Study; HD-YAS, HD Young Adult Study; MPRAGE, Magnetization Prepared Rapid Gradient Echo; ST, slice thickness; T, tesla; T1W, T1 weighted; TE, echo time; TI, inversion time; TR, repetition time.

*Supplementary Table 2: Visual quality control severity rating scale for raw T1-weighted MRI*

| Severity | Rating | Artefact density | Distribution through scan | Image disruption |
|---|---|---|---|---|
| **None** | **0** | No artefacts detected | Throughout the image | None |
| **Mild** | **1** | Minor artefacts | On many slices throughout the image | Small distortion of anatomical structures |
| | **2** | Slightly more prominent | Restricted to one or two slices | |
| **Moderate** | **3** | Slightly more prominent | On many slices throughout the image | Slight distortion of anatomical structures |
| **Severe** | **4** | Heavy artefacts | Restricted to one or two slices | Significantly distort anatomical structures |
| | **5** | Heavy artefacts | Throughout the image in both hemispheres, distorting anatomical structures | |

Raw T1W MRI with scores >3 excluded. QC, quality control; T1W, T1-weighted.

*Supplementary Table 3: Pseudo-pair demographic information for both pipelines*

| Pseudo-pair | Sex | Age | |
|---|---|---|---|
| | | Mean (SD) | Range |
| **Pipeline 1** | | | |
| 1 | F | 39.27 (2.13) | 38.1 – 43.6 |
| 2 | F | 44.95 (0.41) | 44.2 – 45.4 |
| 3 | M | 38.57 (1.16) | 37.2 – 39.8 |
| 4 | M | 40.93 (1.06 | 39.6 – 42.7 |
| **Pipeline 2** | | | |
| 1 | F | 34.72 (2.52) | 30.8 – 38.4 |
| 2 | F | 37.65 (3.21) | 35.2 – 43.6 |
| 3 | F | 39.67 (2.69) | 38.1 – 45.1 |
| 4 | F | 41.58 (2.14) | 39.8 – 42.5 |
| 5 | F | 44.63 (1.59) | 41.8 – 46.6 |
| 6 | F | 45.93 (1.89) | 43.5 – 49.1 |
| 7 | F | 49.92 (2.87) | 44.2 – 51.9 |
| 8 | F | 52.08 (3.67) | 44.9 – 55.1 |
| 9 | M | 35.87 (1.93) | 33.8 – 39.5 |
| 10 | M | 37.52 (1.36) | 35.4 – 39.5 |
| 11 | M | 38.33 (1.14) | 37.0 – 39.9 |
| 12 | M | 40.50 (1.17) | 39.6 – 42.7 |
| 13 | M | 43.65 (3.85) | 40.4 – 50.7 |
| 14 | M | 45.67 (3.68) | 41.8 – 52.6 |
| 15 | M | 47.17 (3.70) | 43.0 – 54.1 |
| 16 | M | 48.72 (3.35) | 43.8 – 54.2 |

Age is reported as mean (SD) and range across all scanners within each pseudo-pair. Each pseudo-pair consists of one healthy control per scanner matched on age and sex. F, female; M, male; SD, standard deviation.

*Supplementary Table 4: RMD participants used per scanner for pipeline optimization and harmonization assessment*

| Study | Healthy controls (n) | pre-CMD (n) | post-CMD (n) | Total (n) |
|---|---|---|---|---|
| **HD-YAS London** | 16 | 16 | - | 32 |
| **HD-CSF London** | 16 | 16 | 20 | 52 |
| **TRACK-HD London** | 4 | 4 | 7 | 15 |
| **TRACK-HD Leiden** | 4 | 4 | 7 | 15 |
| **TRACK-HD Paris** | 4 | 4 | 7 | 15 |
| **TRACK-HD Vancouver** | 4 | 4 | 7 | 15 |
| **Total** | **48** | **48** | **48** | **144** |

Each TRACK-HD site operated an independent scanner; HD-YAS and HD-CSF were each acquired on a single scanner. HD, Huntington's disease; HD-CSF, HD Cerebrospinal Fluid Study; HD-YAS, HD Young Adult Study; post-CMD, pwHD after clinical motor diagnosis; pre-CMD, pwHD before clinical motor diagnosis.

*Supplementary Table 5: FMD participants for scaled-up harmonization.*

| Study | Healthy controls (n) | pre-CMD (n) | post-CMD (n) | Total (n) |
|---|---|---|---|---|
| **HD-YAS London** | 61 | 62 | - | 123 |
| **HD-CSF London** | 15 | 16 | 33 | 64 |
| **TRACK-HD London** | 29 | 30 | 30 | 89 |
| **TRACK-HD Leiden** | 30 | 30 | 30 | 90 |
| **TRACK-HD Paris** | 30 | 30 | 26 | 87 |
| **TRACK-HD Vancouver** | 33 | 30 | 32 | 95 |
| **Total** | **198** | **198** | **151** | **548** |

Each TRACK-HD site operated an independent scanner; HD-YAS and HD-CSF were each acquired on a single scanner. HD, Huntington's disease; HD-CSF, HD Cerebrospinal Fluid Study; HD-YAS, HD Young Adult Study; post-CMD, pwHD after clinical motor diagnosis; pre-CMD, pwHD before clinical motor diagnosis.

*Supplementary Table 6: Visual quality control severity rating scale for grey and white matter tissue probability maps.*

| Severity | Rating | Artefact density | Distribution through scan | Image disruption |
|---|---|---|---|---|
| **0** | **None** | No segmentation errors detected | Throughout the image | None |
| **1** | **Mild** | Minor misclassifications of boundary errors | On many slices throughout the image | Small tissue class misclassification |
| **2** | | Slightly more prominent | Restricted to one or two slices | |
| **3** | **Moderate** | Slightly noticeable boundary inaccuracies or tissue misclassification | On many slices throughout the image | Slight incorrect tissue classification |
| **4** | **Severe** | Heavy artefacts | Restricted to one or two slices | Significantly distort anatomical structures |
| **5** | | Heavy artefacts | Throughout the image in both hemispheres, distorting anatomical structures | |

TPMs with scores >3 excluded. If the TPM for one tissue class was excluded, all corresponding TPMs for that participant were excluded, both unharmonized and harmonized. QC, quality control; TPM, tissue probability map.

*Supplementary Table 7: Criteria for go/no-go decisions for pipeline selection.*

| Category | Assessment method from each pipeline | Outcome measures to compare between pipelines | Deciding factors to choose a pipeline for FMD |
|---|---|---|---|
| | | **Image quality assessments** | |
| **Visual QC** | T1W MRI visual QC | Artifact rating scale (0-5). | Low average artifact rating, with a minimal number of scans rated 4 or higher. |
| **Segmentation consistency** | Grey/white matter TPM overlay | Visual comparison of segmentation overlays. | Minimal significant distortion of anatomical boundaries and may even demonstrate improved tissue assignment. |
| **Image quality metrics** | MRIQC (SNR, CNR) | SNR and CNR values. | SNR and CNR values that are stable or higher than the comparative pipeline. |
| **Voxel contrast intensity distributions** | Intensity plots | Plots of intensity distributions from different scanners. | More aligned intensity distributions from different scanners while preserving biologically meaningful variation (tissue-class and disease-related differences). |
| | | **Voxel-wise assessments** | |
| **Scanner-wise assessment** | Paired voxel-wise analysis (flexible factorial GLM) | Peak T-values; number of significant voxels/clusters; spatial location of clusters. | Fewer/smaller scanner-driven clusters, lower peak T-values, minimal overlap with biologically important ROIs. |
| **Within disease groups** | Paired voxel-wise analysis (flexible factorial GLM) | Peak T-values; number of significant voxels/clusters; spatial location of clusters. | Fewer/smaller within-group clusters, lower peak T-values, minimal overlap with biologically important ROIs. |
| **Disease group preservation** | Independent voxel-wise analysis (ANOVA) | Peak T-values; number of significant voxels/clusters; spatial location of clusters. | Disease-related clusters, higher peak T-values, spatial location within expected disease-relevant ROIs. |
| **Relationships with clinical measures** | Independent voxel-wise analysis (flexible factorial GLM) | Peak T-values; number of significant voxels/clusters; spatial location of clusters. | Disease-related clusters, higher peak T-values, spatial location within expected disease-relevant ROIs. |
| | | **Volumetric assessments** | |
| **Overall volume changes** | Linear mixed-effects modelling of volumes from TPMs | Harmonization main effect (β, 95% CI, p-value). Post-hoc pairwise contrasts (Bonferroni-adjusted p-values). Mean differences (harmonized − unharmonized). Model fit comparison (ΔAIC/ΔBIC). | Minimal systematic bias (small % bias), non-significant or very small harmonization effect size, and no deterioration in model fit. |
| **Scanner/disease-specific biases** | Linear mixed-effects modelling of volumes from TPMs | Interaction effects (harmonization × scanner; harmonization × disease: β, 95% CI, p-value). Scanner-/disease-wise post-hoc contrasts (Bonferroni-adjusted p-values). Mean differences by scanner/disease group. Model fit comparison (ΔAIC/ΔBIC). | Reduced scanner-driven differences (small/non-significant interactions, small effect sizes), no new scanner × disease interactions, and mean differences within acceptable thresholds. |
| **Agreement across pipelines** | ICC, spaghetti plots, Bland-Altman plots | ICC values (0–1), visual inspection of spaghetti and Bland-Altman plots for systematic bias. | High ICC (≥0.75 indicating good agreement); spaghetti/Bland-Altman plots showing minimal systematic bias in volumes after harmonization. |
| | | **Advanced assessments** | |
| **Principal Component Analysis (PCA)** | PCA on Voxel Intensities | Variance explained by top components; clustering by scanner/disease | Less scanner-driven clustering, lower variance explained by scanner-related PCs |
| **SP-ComBat re-estimation** | Re-estimation of parametric maps | Gamma and delta intensity maps | Additive terms close to zero and multiplicative terms close to one, indicating minimal scanner-driven deviation and alignment to a unified standard distribution. |
| **CNN classifiers** | CNN for scanner prediction (FMD only) | Scanner prediction accuracy | Lower accuracy (closer to chance), meaning stronger harmonization |

AIC, Akaike information criterion; ANOVA, analysis of variance; BIC, Bayesian information criterion; CNN, convolutional neural network; CNR, contrast to noise ratio; FMD, full multi-study dataset; GLM, general linear model; MRI, magnetic resonance imaging; PCA, Principal Component Analysis; PC, Principal Component; ROI, region of interest; SNR, signal to noise ratio; SP, superpixel; TPM, tissue probability map; QC, quality control.

*Supplementary Table 8: Specification of F- and T-contrast vectors for voxel-wise harmonization and disease group analyses in grey and white matter.*

| Comparison | Contrast | Vector |
|---|---|---|
| | Within scanner-specific contrasts | |
| Main effect of harmonization | F-contrast | [1 -1 1 -1 1 -1 1 -1 1 -1 1 -1] |
| Harmonized > Unharmonized across all scanners | T-contrast | [-1 1 -1 1 -1 1 -1 1 -1 1 -1 1] |
| Harmonized > Unharmonized THD Paris | T-contrast | [-1 0 0 0 0 0 1 0 0 0 0 0] |
| Harmonized > Unharmonized THD London | T-contrast | [0 -1 0 0 0 0 0 1 0 0 0 0] |
| Harmonized > Unharmonized THD Leiden | T-contrast | [0 0 -1 0 0 0 0 0 1 0 0 0] |
| Harmonized > Unharmonized THD Vancouver | T-contrast | [0 0 0 -1 0 0 0 0 0 1 0 0] |
| Harmonized > Unharmonized HD-YAS | T-contrast | [0 0 0 0 -1 0 0 0 0 0 1 0] |
| Harmonized > Unharmonized HD-CSF | T-contrast | [0 0 0 0 0 -1 0 0 0 0 0 1] |
| | Within disease group specific contrasts | |
| Main effect of harmonization | F-contrast | [1 -1 1 -1 1 -1] |
| Harmonized > Unharmonized across all disease groups | T-contrast | [-1 1 -1 1 -1 1] |
| Harmonized > Unharmonized HC | T-contrast | [0 -1 0 0 1 0] |
| Harmonized > Unharmonized pre-CMD | T-contrast | [-1 0 0 1 0 0] |
| Harmonized > Unharmonized post-CMD | T-contrast | [0 0 -1 0 0 1] |
| | Between disease group specific contrasts | |
| HCs > pre-CMD | T-contrast | [1 -1 0] |
| HCs > post-CMD | T-contrast | [1 0 -1] |

[1] represents unharmonized and [-1] represents harmonized. [0] was used to block comparisons in the relevant scanners or disease groups. HC, healthy control; HD, Huntington's disease; HD-CSF, HD Cerebrospinal Fluid Study; HD-YAS, HD Young Adult Study; post-CMD, pwHD after clinical motor diagnosis; pre-CMD, pwHD before clinical motor diagnosis; THD, TRACK-HD study site.

*Supplementary Table 9: RMD group demographics*

| Study | Disease group (n) | Age | | Sex | | CAG | | DBS | |
|---|---|---|---|---|---|---|---|---|---|
| | | mean (SD), range | p | %M | p | mean (SD), range | p | mean (SD), range | p |
| **HD-YAS London** | Control (16) | 36.5 (2.1), 34 – 41.5 | 0.145[1] | 56 | 1.000 | - | - | - | - |
| | pre-CMD (16) | 35.4 (2), 32 - 40 | | 56 | | 41.2 (0.9), 40 - 43 | | 200 (27.6), 153 - 240 | |
| **HD-CSF London** | Control (16) | 50.4 (11), 29 - 74 | | 56 | | - | | - | |
| | pre-CMD (16) | 41.5 (11.7), 25 - 67 | 0.026[1], 0.194[2], < 0.005[3] | 56 | 0.9078 | 42.3 (1.7), 40 - 45 | 0.657 | 272.2 (66.3), 113 - 383 | < 0.005 |
| | post-CMD (20) | 54.6 (8.8), 4.9 – 71.5 | | 50 | | 42.5 (1.7), 40 - 45 | | 370 (72.7), 248 - 493 | |
| **TRACK-HD London** | Control (4) | 50.8 (8.9), 42.6 – 63.5 | | 50 | | - | | - | |
| | pre-CMD (4) | 36.8 (3.7), 33.6 – 41 | 0.033[1], 0.810[2], 0.012[3] | 0 | 0.1580 | 42.8 (1.5), 41 - 44 | 0.648 | 263.3 (34.5), 213 - 287 | 0.015 |
| | post-CMD (7) | 52.1 (9.4), 35.8 – 63.8 | | 43 | | 42.3 (1.6), 40 - 45 | | 342.3 (45.1), 287 - 406 | |
| **TRACK-HD Leiden** | Control (4) | 52.3 (5.3), 47.3 – 58.2 | | 50 | | - | | - | |
| | pre-CMD (4) | 36.2 (0.7), 36.7 – 36.9 | 0.017[1], 0.97[2], 0.008[3] | 25 | 0.7059 | 43.8 (0.96), 43 - 45 | 0.320 | 298 (30.8), 270 - 336 | 0.008 |
| | post-CMD (7) | 52.5 (11), 31.7 – 63.9 | | 29 | | 42.7 (1.8), 41 - 46 | | 362.2 (29.7), 321 - 398 | |
| **TRACK-HD Paris** | Control (4) | 48 (8.2), 38.6 – 58.5 | | 50 | | - | | - | |
| | pre-CMD (4) | 35.3 (2.7), 32.4 – 37.8 | 0.031[1], 0.343[2], 0.002[3] | 25 | 0.7537 | 43 (1.4), 42 - 45 | 0.655 | 263.2 (39.1), 218 - 308 | 0.002 |
| | post-CMD (7) | 52.6 (8.4), 37.4 – 60.9 | | 43 | | 42.6 (1.5), 42 - 46 | | 363 (35.3), 313 - 396 | |
| **TRACK-HD Vancouver** | Control (4) | 51.4 (5.5), 45.6 – 58.1 | | 50 | | - | | - | |
| | pre-CMD (4) | 36.2 (8.4), 27.5 – 47.3 | 0.029[1], 0.895[2], 0.012[3] | 75 | 0.7537 | 43 (1.4), 41 - 44 | 0.902 | 262.9 (21.4), 234 - 280 | < 0.005 |
| | post-CMD (7) | 52.1 (9.9), 31.1 – 60.2 | | 57 | | 42.9 (2), 41 - 47 | | 367 (21.8), 331 - 391 | |

Data are expressed as mean (SD), range. Disease groups were compared using Independent One-Way ANOVA (Age, CAG and DBS) and Chi-squared test (Sex). For age, post-estimation Wald tests were used to assess intergroup differences with Bonferroni correction for multiple comparisons. Bold values represent significant comparisons. [1]HC vs pre-CMD, [2]HC vs post-CMD, and [3]pre-CMD vs post-CMD. CAG, cytosine-adenine-guanine; DBS, disease burden score; HC, healthy control; HD, Huntington's disease; HD-CSF, HD Cerebrospinal Fluid Study; HD-YAS, HD Young Adult Study; M, male; SD, standard deviation; post-CMD, pwHD after clinical motor diagnosis; pre-CMD, pwHD before clinical motor diagnosis.

*Supplementary Table 10: FMD group demographics*

| Study | Disease group (n) | Age | | Sex | | CAG | | DBS | |
|---|---|---|---|---|---|---|---|---|---|
| | | mean (SD), range | p | %M | p | mean (SD), range | p | mean (SD), range | p |
| **HD-YAS London** | Control (61) | 29.7 (5.5), 20.2 – 40 | 0.941[1] | 39 | 0.4054 | - | - | - | - |
| | pre-CMD (62) | 29.6 (5.7), 19.3 – 40.9 | | 47 | | 42.3 (1.7), 39 - 47 | | 196.2 (37.2), 122 - 316 | |
| **HD-CSF London** | Control (15) | 49.5 (11), 30 – 74.7 | | 53 | | - | | - | |
| | pre-CMD (16) | 42 (11.7), 26 – 67.5 | 0.050[1], 0.116[2], < 0.005[3] | 56 | 0.8672 | 42.3 (1.7), 40 - 45 | 0.474 | 284 (68.2), 122 – 398 | < 0.005 |
| | post-CMD (27) | 54.8 (9.2), 40.7 – 71.5 | | 48 | | 42.7 (1.9), 39 - 45 | | 396.8 (103), 191 - 622 | |
| **TRACK-HD London** | Control (29) | 45.1 (9.2), 24 – 63.5 | | 45 | | - | | - | |
| | pre-CMD (30) | 40.8 (7.5), 26.1 – 58.5 | 0.069[1], 0.150[2], 0.001[3] | 47 | 0.8657 | 43.1 (1.9), 40 – 48 | 0.478 | 305.8 (43.7), 221 – 386 | < 0.005 |
| | post-CMD (30) | 48.4 (9.8), 29.6 – 64.1 | | 40 | | 43.5 (2.4), 39 - 48 | | 378.4 (79.5), 232 - 541 | |
| **TRACK-HD Leiden** | Control (30) | 49.2 (8.3), 35.2 – 65.7 | | 47 | | - | | - | |
| | pre-CMD (30) | 43.8 (8), 26.4 – 61.7 | 0.023[1], 0.821[2], 0.044[3] | 40 | 0.5283 | 42.5 (2.4), 39 – 50 | 0.158 | 301.4 (53.7), 181 - 399 | < 0.005 |
| | post-CMD (28) | 48.6 (10.5), 30.7 – 63.9 | | 32 | | 43.4 (2.5), 40 - 50 | | 374.1 (56.4), 284 - 490 | |
| **TRACK-HD Paris** | Control (29) | 44.4 (11.2), 25.9 – 63 | | 48 | | - | | - | |
| | pre-CMD (30) | 39.1 (9.1), 22.3 – 64.1 | 0.036[1], 0.064[2], < 0.005[3] | 53 | 0.6346 | 43.7 (2.7) 39 - 52 | 0.299 | 308 (49.7), 207 - 415 | 0.001 |
| | post-CMD (27) | 49.3 (8.4), 33.7 – 60.9 | | 41 | | 43 (2), 39 - 47 | | 368.8 (74.4), 165 - 507 | |
| **TRACK-HD Vancouver** | Control (33) | 45.7 (12), 23 – 62.6 | | 39 | | - | | - | |
| | pre-CMD (30) | 39.6 (10.3), 18.6 – 61.8 | 0.035[1], 0.371[2], 0.004[3] | 40 | 0.007 | 43.3 (2.5), 40 - 50 | 0.135 | 296.8 (43.1), 219 - 373 | < 0.005 |
| | post-CMD (31) | 48.2 (11.3), 22.8 – 63.4 | | 74 | | 44.7 (4.2), 40 - 59 | | 412.3 (75.2), 280 - 574 | |

Data are expressed as mean (SD), range. Disease groups were compared using Independent One-Way ANOVA (Age, CAG and DBS) and Chi-squared test (Sex). For age, post-estimation Wald tests were used to assess intergroup differences with Bonferroni correction for multiple comparisons. Bold values represent significant comparisons. [1]HC vs pre-CMD, [2]HC vs post-CMD and [3]pre-CMD vs post-CMD. CAG, cytosine-adenine-guanine; DBS, disease burden score; HC, healthy control; HD, Huntington's disease; HD-CSF, HD Cerebrospinal Fluid Study; HD-YAS, HD Young Adult Study; M, male; SD, standard deviation; post-CMD, pwHD after clinical motor diagnosis; pre-CMD, pwHD before clinical motor diagnosis.

*Supplementary Table 11: MRIQC image quality metric differences between unharmonized and harmonized T1W MRI.*

| | Adjusted MD | SE | 95 % CI | p-value |
|---|---|---|---|---|
| | | **CNR** | | |
| Overall | -0.018 | 0.037 | -0.090 to 0.055 | 0.632 |
| HD-YAS | 0.681 | 0.077 | 0.531 to 0.832 | **< 0.0001** |
| HD-CSF | 0.247 | 0.113 | 0.027 to 0.468 | 0.028 |
| THD London | -0.052 | 0.091 | -0.230 to 0.127 | 0.570 |
| THD Leiden | -0.505 | 0.091 | -0.684 to -0.326 | **< 0.0001** |
| THD Paris | 0.008 | 0.092 | -0.173 to 0.190 | 0.929 |
| THD Vancouver | -0.639 | 0.088 | -0.813 to -0.466 | **< 0.0001** |
| | | **CJV** | | |
| Overall | -0.084 | 0.004 | -0.092 to -0.077 | **< 0.0001** |
| HD-YAS | -0.181 | 0.008 | -0.197 to -0.166 | **< 0.0001** |
| HD-CSF | -0.119 | 0.012 | -0.141 to -0.096 | **< 0.0001** |
| THD London | -0.058 | 0.009 | -0.077 to -0.040 | **< 0.0001** |
| THD Leiden | -0.016 | 0.009 | -0.034 to 0.003 | 0.098 |
| THD Paris | -0.075 | 0.010 | -0.094 to -0.057 | **< 0.0001** |
| THD Vancouver | -0.033 | 0.009 | -0.051 to -0.015 | **< 0.0001** |
| | | **SNR (total)** | | |
| Overall | 4.449 | 0.060 | 4.332 to 4.566 | **< 0.0001** |
| HD-YAS | 8.541 | 0.124 | 8.298 to 8.785 | **< 0.0001** |
| HD-CSF | 5.450 | 0.182 | 5.094 to 5.806 | **< 0.0001** |
| THD London | 2.279 | 0.147 | 1.991 to 2.566 | **< 0.0001** |
| THD Leiden | 3.124 | 0.147 | 2.835 to 3.413 | **< 0.0001** |
| THD Paris | 2.311 | 0.149 | 2.019 to 2.604 | **< 0.0001** |
| THD Vancouver | 3.684 | 0.143 | 3.404 to 3.964 | **< 0.0001** |
| | | **SNR (GM)** | | |
| Overall | 1.650 | 0.047 | 1.558 to 1.741 | **< 0.0001** |
| HD-YAS | 4.223 | 0.097 | 4.033 to 4.414 | **< 0.0001** |
| HD-CSF | 3.531 | 0.142 | 3.252 to 3.809 | **< 0.0001** |
| THD London | -0.606 | 0.115 | -0.831 to -0.381 | **< 0.0001** |
| THD Leiden | 1.318 | 0.115 | 1.092 to 1.544 | **< 0.0001** |
| THD Paris | -0.851 | 0.117 | -1.080 to -0.623 | **< 0.0001** |
| THD Vancouver | 1.829 | 0.112 | 1.610 to 2.048 | **< 0.0001** |
| | | **SNR (WM)** | | |
| Overall | 9.751 | 0.140 | 9.476 to 10.026 | **< 0.0001** |
| HD-YAS | 19.376 | 0.292 | 18.802 to 19.949 | **< 0.0001** |
| HD-CSF | 11.067 | 0.428 | 10.229 to 11.905 | **< 0.0001** |
| THD London | 5.898 | 0.345 | 5.222 to 6.575 | **< 0.0001** |
| THD Leiden | 5.621 | 0.347 | 4.940 to 6.301 | **< 0.0001** |
| THD Paris | 6.010 | 0.351 | 5.322 to 6.699 | **< 0.0001** |
| THD Vancouver | 7.179 | 0.336 | 6.520 to 7.837 | **< 0.0001** |
| | | **SNR (CSF)** | | |
| Overall | 1.946 | 0.035 | 1.878 to 2.014 | **< 0.0001** |
| HD-YAS | 2.024 | 0.073 | 1.882 to 2.167 | **< 0.0001** |
| HD-CSF | 1.753 | 0.106 | 1.545 to 1.961 | **< 0.0001** |
| THD London | 1.544 | 0.086 | 1.376 to 1.712 | **< 0.0001** |
| THD Leiden | 2.433 | 0.086 | 2.264 to 2.601 | **< 0.0001** |
| THD Paris | 1.775 | 0.087 | 1.604 to 1.946 | **< 0.0001** |
| THD Vancouver | 2.044 | 0.083 | 1.881 to 2.207 | **< 0.0001** |

Data are derived from linear mixed-effects models between unharmonized and harmonized metrics; adjusted MD represents the average marginal effect. CNR is defined between GM and WM. CJV reflect field inhomogeneity. The definition of noise in SNR is the within-tissue variance. CI, confidence interval; CJV, coefficient of joint variation; CNR, contrast-to-noise ratio; CSF, cerebrospinal fluid; GM, grey matter; HD, Huntington's disease; HD-CSF, HD Cerebrospinal Fluid Study; HD-YAS, HD Young Adult Study; MD, mean difference; SE, standard error; SNR, signal-to-noise ratio; THD, TRACK-HD study site; WM, white matter.

*Supplementary Table 12: Variance explained by Principal Components in unharmonized and harmonized T1W MRI*

| PCA Component | Variance explained by Principal Components (%) | |
|---|---|---|
| | Unharmonized | Harmonized |
| **PC1** | 97.61 | 35.68 |
| **PC2** | 0.59 | 15.78 |
| **PC3** | 0.41 | 10.02 |
| **PC4** | 0.32 | 8.95 |

PC, principal component; PCA, principal component analysis.

*Supplementary Table 13: Variance explained by Principal Components in unharmonized and harmonized healthy control T1W MRI only*

| PCA Component | Variance explained by Principal Components (%) | |
|---|---|---|
| | Unharmonized | Harmonized |
| **PC1** | 96.84 | 31.14 |
| **PC2** | 0.77 | 15.50 |
| **PC3** | 0.52 | 11.10 |
| **PC4** | 0.38 | 9.22 |

PC, principal component; PCA, principal component analysis.

*Supplementary Table 14: QC outcomes for grey and white matter TPMs, both before and after harmonization.*

| Study | Unharmonized | | | Harmonized | | |
|---|---|---|---|---|---|---|
| | Scores 0-2 (n) | Score 3-5 (n) | | Scores 0-2 (n) | Score 3-5 (n) | |
| | | Non-systematic | Systematic | | Non-systematic | Systematic |
| | | | Grey matter | | | |
| **HD-YAS London** | 3 | 3 | 117 | 120 | 3 | 0 |
| **HD-CSF London** | 45 | 10 | 3 | 49 | 9 | 0 |
| **TRACK-HD London** | 85 | 4 | 0 | 87 | 2 | 0 |
| **TRACK-HD Leiden** | 81 | 7 | 0 | 81 | 7 | 0 |
| **TRACK-HD Paris** | 83 | 3 | 0 | 83 | 3 | 0 |
| **TRACK-HD Vancouver** | 87 | 7 | 0 | 85 | 9 | 0 |
| **Total** | 384 | 34 | 120 | 505 | 33 | 0 |
| | | | White matter | | | |
| **HD-YAS London** | 6 | 0 | 117 | 123 | 0 | 0 |
| **HD-CSF London** | 55 | 0 | 3 | 58 | 0 | 0 |
| **TRACK-HD London** | 87 | 2 | 0 | 89 | 0 | 0 |
| **TRACK-HD Leiden** | 88 | 0 | 0 | 88 | 0 | 0 |
| **TRACK-HD Paris** | 84 | 2 | 0 | 84 | 2 | 0 |
| **TRACK-HD Vancouver** | 94 | 0 | 0 | 94 | 0 | 0 |
| **Total** | 414 | 4 | 120 | 536 | 2 | 0 |

Segmentation QC ratings before and after harmonization by scanner, shown separately for grey matter and white matter TPMs before and after harmonization. Systematic errors refer to consistent, scanner-driven misclassification affecting scans from a given scanner, including grey matter over-segmentation and white matter under-segmentation; non-systematic errors refer to sporadic tissue misclassifications unrelated to scanner identity, most commonly meningeal over-segmentation. Grey and white matter, and unharmonized and harmonized counts are independent, as a participant may pass QC at one stage but not the other; the 501 participants taken forward for analysis represent those passing QC in both statuses across both tissue classes. Where either tissue class failed QC, all corresponding TPMs were excluded for that participant across both harmonization statuses. Scans scoring >3 due to systematic errors were retained in harmonization pipeline comparisons to characterize the full extent of segmentation failure and its resolution; all scans scoring >3 were excluded from biological preservation analyses (Section 3.4). HD, Huntington's disease; HD-CSF, HD Cerebrospinal Fluid Study; HD-YAS, HD Young Adult Study; QC, quality control; TPM, tissue probability map.

*Supplementary Table 15: Scanner specific voxel-wise statistics between unharmonized and harmonized grey and white matter TPMs*

| Contrast (F or T contrast) | Total clusters | Largest cluster | Peak score | MNI coordinate | Atlas Location | p |
|---|---|---|---|---|---|---|
| | *N* | *n voxels* | *F/T-score* | *x, y, z (mm)* | *Grey matter: AAL3 White matter: HCPEx* | *FWE-corrected* |
| **Grey matter** | | | | | | |
| **Main effect of harmonization (F)** | 182 | 387 | 281.5 | 2, 26, -26 | R rectus | < 0.0001 |
| **Overall (T)** | 88 | 518 | 13.5 | -12, -30, -2 | L Thalamus | < 0.0001 |
| **THD London (T)** | 29 | 2647 | 12.9 | -14, -4, -32 | L parahippocampal gyrus | < 0.0001 |
| **THD Leiden (T)** | 21 | 1214 | 13.5 | -3, -8, -14 | L substantia nigra | < 0.0001 |
| **THD Paris (T)** | 33 | 4334 | 11.32 | 8, -2, -15 | R substantia nigra | < 0.0001 |
| **THD Vancouver (T)** | 12 | 759 | 10.36 | 8, 12, -22 | R rectus | < 0.0001 |
| **HD-YAS (T)** | 84 | 6491 | 25.93 | 8, -12, 15 | R thalamus | < 0.0001 |
| **HD-CSF (T)** | 16 | 33 | 6.1 | 27, -58, -34 | R cerebellum | < 0.0001 |
| **White matter** | | | | | | |
| **Main effect of harmonization (F)** | 68 | 3110 | 181 | -14, -18, 18 | L internal capsule | < 0.0001 |
| **Overall (T)** | 14 | 392 | 11.6 | 16, 20, -16 | Undefined | < 0.0001 |
| **THD London (T)** | 12 | 112 | 8.08 | 6, -28, -4 | Undefined | < 0.0001 |
| **THD Leiden (T)** | 14 | 66 | 8.88 | 32, -2, 4 | R internal capsule | < 0.0001 |
| **THD Paris (T)** | 15 | 29 | 7.4 | 8, -68, -28 | Undefined | < 0.0001 |
| **THD Vancouver (T)** | 9 | 219 | 10.99 | -30, -2, 8 | L internal capsule | < 0.0001 |
| **HD-YAS (T)** | 16 | 4582 | 21.09 | 14, 15, 54 | Undefined | < 0.0001 |
| **HD-CSF (T)** | 32 | 56 | 10.35 | 6, -54, -18 | Undefined | < 0.0001 |

Summary statistics are presented for significant clusters and peaks from grey matter and white matter scanner-wise voxel-based comparisons. Peak scores are from the largest significant cluster with the corresponding MNI coordinate. T contrasts are harmonized > unharmonized. Z-scores were omitted because all largest T- and F-values exceeded the finite range of the T/F→Z transformation, producing Z = Inf. Peak statistics are therefore reported using the original T- or F-values together with FWE-corrected p-values. AAL3, Automated Anatomical Labelling atlas 3; FWE, family-wise error; HCPEx, Extended Human Connectome Project multimodal parcellation atlas; HD, Huntington's disease; HD-CSF, HD Cerebrospinal Fluid Study; HD-YAS, HD Young Adult Study; L, left; MNI, Montreal Neurological Institute; R, right; THD, TRACK-HD study site.

*Supplementary Table 16: Scanner specific volumetric differences between unharmonized and harmonized grey and white matter.*

| | Tissue | Adjusted MD | SE | 95% CI | p-value |
|---|---|---|---|---|---|
| **Overall volumetric differences** | | | | | |
| **Overall** | Grey matter | 11.878 | 1.323 | 9.286 to 14.471 | < 0.0001 |
| | White matter | -17.623 | 0.837 | -19.263 to -15.983 | < 0.0001 |
| **Scanner specific volumetric differences** | | | | | |
| **HD-YAS** | Grey matter | 13.245 | 1.235 | 9.988 to 16.502 | < 0.0001 |
| | White matter | -15.554 | 0.781 | -17.614 to -13.494 | < 0.0001 |
| **HD-CSF** | Grey matter | -38.778 | 1.823 | -43.588 to -33.967 | < 0.0001 |
| | White matter | 18.425 | 1.153 | 15.382 to 21.467 | < 0.0001 |
| **TRACK-HD London** | Grey matter | 23.99 | 1.362 | 20.397 to 27.583 | < 0.0001 |
| | White matter | -17.513 | 0.861 | -19.785 to -15.24 | < 0.0001 |
| **TRACK-HD Leiden** | Grey matter | 12.429 | 1.395 | 8.748 to 16.11 | < 0.0001 |
| | White matter | -12.256 | 0.882 | -14.584 to -9.928 | < 0.0001 |
| **TRACK-HD Paris** | Grey matter | 24.799 | 1.387 | 21.139 to 28.459 | < 0.0001 |
| | White matter | -16.832 | 0.877 | -19.146 to -14.518 | < 0.0001 |
| **TRACK-HD Vancouver** | Grey matter | 7.895 | 1.364 | 4.296 to 11.495 | < 0.0001 |
| | White matter | -14.146 | 0.863 | -16.422 to -11.87 | < 0.0001 |

Values obtained from post-hoc pairwise comparisons of estimated marginal means (Bonferroni-corrected) to quantify overall and scanner-specific differences between unharmonized and harmonized raw grey and white matter volumes. CI, confidence interval; HD, Huntington's disease; HD-CSF, HD Cerebrospinal Fluid Study; HD-YAS, HD Young Adult Study; MD, mean difference; SE, standard error.

*Supplementary Table 17: Within disease-specific voxel-wise statistics between unharmonized and harmonized grey and white matter TPMs*

| Contrast (F or T contrast) | Total clusters | Largest cluster | Peak score | MNI coordinate | Atlas Location | p |
|---|---|---|---|---|---|---|
| | *N* | *n voxels* | *F/T-score* | *x, y, z (mm)* | *Grey matter: AAL3 White matter: HCPEx* | *FWE-corrected* |
| **Grey matter** | | | | | | |
| **Main effect of harmonization (F)** | 14 | 162 | 88.89 | -32, -4, 3 | L putamen | < 0.0001 |
| **Overall (T)** | 14 | 589 | 7.27 | 6, 2, -10 | R nuclear accumbens | < 0.0001 |
| **Healthy control (T)** | 26 | 3053 | 11.65 | 6, 2, -10 | R nuclear accumbens | < 0.0001 |
| **pre-CMD (T)** | 28 | 1992 | 10.51 | 8, 2, -10 | R nuclear accumbens | < 0.0001 |
| **post-CMD (T)** | 24 | 743 | 9.19 | 8, 2, -10 | R nuclear accumbens | < 0.0001 |
| **White matter** | | | | | | |
| **Main effect of harmonization (F)** | 37 | 290 | 108.4 | 21, 16, 2 | R internal capsule | < 0.0001 |
| **Overall (T)** | 5 | 120 | 9.33 | -32, -3, 8 | L internal capsule | < 0.0001 |
| **Healthy control (T)** | 13 | 201 | 13.2 | -32, -3, 8 | L internal capsule | < 0.0001 |
| **pre-CMD (T)** | 13 | 132 | 11.08 | -32, -3, 8 | L internal capsule | < 0.0001 |
| **post-CMD (T)** | 23 | 42 | 9.05 | 4, -54, -18 | Undefined | < 0.0001 |

Summary statistics are presented for significant clusters and peaks from grey matter and white matter within-disease voxel-based comparisons with all those QC >3 removed. Peak scores are from the largest significant cluster with the corresponding MNI coordinate. T contrasts are harmonized > unharmonized. Z-scores were omitted because all largest T- and F-values exceeded the finite range of the T/F→Z transformation, producing Z = Inf. Peak statistics are therefore reported using the original T- or F-values together with FWE-corrected p-values. AAL3, Automated Anatomical Labelling atlas 3; FWE, family-wise error; HCPEx, Extended Human Connectome Project multimodal parcellation atlas; L, left; MNI, Montreal Neurological Institute; post-CMD, pwHD after clinical motor diagnosis; pre-CMD, pwHD before clinical motor diagnosis; QC, quality control; R, right.

*Supplementary Table 18: Within disease specific volumetric differences between unharmonized and harmonized grey and white matter*

| | Tissue | Adjusted MD | SE | 95% CI | p-value |
|---|---|---|---|---|---|
| **Within disease specific volumetric differences** | | | | | |
| **Healthy Controls** | Grey matter | 4.212 | 1.572 | 0.448 to 7.975 | 0.0221 |
| | White matter | -12.569 | 1.049 | -15.081 to -10.057 | < 0.0001 |
| **pre-CMD** | Grey matter | 6.061 | 1.574 | 2.294 to 9.829 | 0.0004 |
| | White matter | -10.842 | 1.05 | -13.357 to -8.327 | < 0.0001 |
| **post-CMD** | Grey matter | 8.424 | 1.61 | 4.57 to 12.278 | < 0.0001 |
| | White matter | -6.75 | 1.075 | -9.323 to -4.177 | < 0.0001 |

Values obtained from post-hoc pairwise comparisons of estimated marginal means (Bonferroni-corrected) from all TPMs QC < 3 to quantify disease-specific differences between unharmonized and harmonized grey and white matter volumes. CI, confidence interval; HD, Huntington's disease; MD, mean difference; post-CMD, pwHD after clinical motor diagnosis; pre-CMD, pwHD before clinical motor diagnosis; QC, quality control; SE, standard error.

*Supplementary Table 19: voxel-wise between disease group differences in unharmonized and harmonized T1W MRI*

| Contrast | Condition | Total clusters | Largest cluster | Peak score | MNI coordinate | Atlas Location | p |
|---|---|---|---|---|---|---|---|
| | | *N* | *n voxels* | *T-score* | *x, y, z (mm)* | *Grey matter: AAL3 White matter: HCPEx* | *FWE-corrected* |
| **Grey matter** | | | | | | | |
| **HC > pre-CMD** | Unharmonized | 18 | 1489 | 11.67 | 30, -10, -8 | R putamen | < 0.0001 |
| | Harmonized | 12 | 3862 | 14.51 | 32, -10, -9 | R putamen | < 0.0001 |
| **HC > post-CMD** | Unharmonized | 167 | 48505 | 24.37 | 22, 10, 2 | R putamen | < 0.0001 |
| | Harmonized | 165 | 39360 | 25.3 | 27, 9, 4 | R putamen | < 0.0001 |
| **White matter** | | | | | | | |
| **HC > pre-CMD** | Unharmonized | 39 | 3706 | 9.03 | -24, -6, -4 | L internal capsule | < 0.0001 |
| | Harmonized | 32 | 3080 | 9.21 | -24, -6, -4 | L internal capsule | < 0.0001 |
| **HC > post-CMD** | Unharmonized | 9 | 102682 | 20.01 | 21, 3, -2 | L internal capsule | < 0.0001 |
| | Harmonized | 6 | 90751 | 20.48 | -21, 0, -2 | L internal capsule | < 0.0001 |

Summary statistics are presented for significant clusters and peaks from grey matter and white matter disease group voxel-based comparisons with all those with < 3 on QC severity scale (N = 379). Peak scores are from the largest significant cluster with the corresponding MNI coordinate. T-contrasts compare healthy controls to pre-CMD and post-CMD participants. Z-scores were omitted because all largest T- and F-values exceeded the finite range of the T/F→Z transformation, producing Z = Inf. Peak statistics are therefore reported using the original T- or F-values together with FWE-corrected p-values. AAL3, Automated Anatomical Labelling atlas 3; FWE, family-wise error; HC, healthy controls; HCPEx, Extended Human Connectome Project multimodal parcellation atlas; L, left; MNI, Montreal Neurological Institute; post-CMD, pwHD after clinical motor diagnosis; pre-CMD, pwHD before clinical motor diagnosis; R, right.

*Supplementary Table 20: Volumetric post-hoc disease group differences in grey and white matter before and after harmonization.*

| Comparison | Condition | Adj. MD | SE | Z | 95% CI | p |
|---|---|---|---|---|---|---|
| **Grey matter** | | | | | | |
| **HC versus pre-CMD** | Unharmonized | -0.22 | 0.36 | -0.63 | -1.11 to 0.66 | 1.000 |
| | Harmonized | -0.1 | 0.36 | -0.27 | -0.98 to 0.79 | 1.000 |
| **HC versus post-CMD** | Unharmonized | -3.93 | 0.36 | -11 | -4.82 to -3.04 | < 0.0001 |
| | Harmonized | -3.61 | 0.36 | -10.1 | -4.5 to -2.72 | < 0.0001 |
| **White matter** | | | | | | |
| **HC versus pre-CMD** | Unharmonized | -0.92 | 0.23 | -4.03 | -1.49 to -0.35 | 0.0002 |
| | Harmonized | -0.81 | 0.23 | -3.53 | -1.38 to -0.24 | 0.0016 |
| **HC versus post-CMD** | Unharmonized | -2.64 | 0.23 | -11.48 | -3.22 to -2.84 | < 0.0001 |
| | Harmonized | -2.27 | 0.23 | -9.85 | -2.84 to -1.69 | < 0.0001 |

Values obtained from post-hoc pairwise comparisons of estimated marginal means (Bonferroni-corrected) to quantify disease-group difference in grey and white volumes before and after harmonization in those with QC severity rating score < 3. Volumes are % of TIV (volume/TIV * 100). CI, confidence interval; HC, healthy control; MD, mean difference; post-CMD, pwHD after clinical motor diagnosis; pre-CMD, pwHD before clinical motor diagnosis; QC, quality control; SE, standard error.

### 4 Supplementary Figures

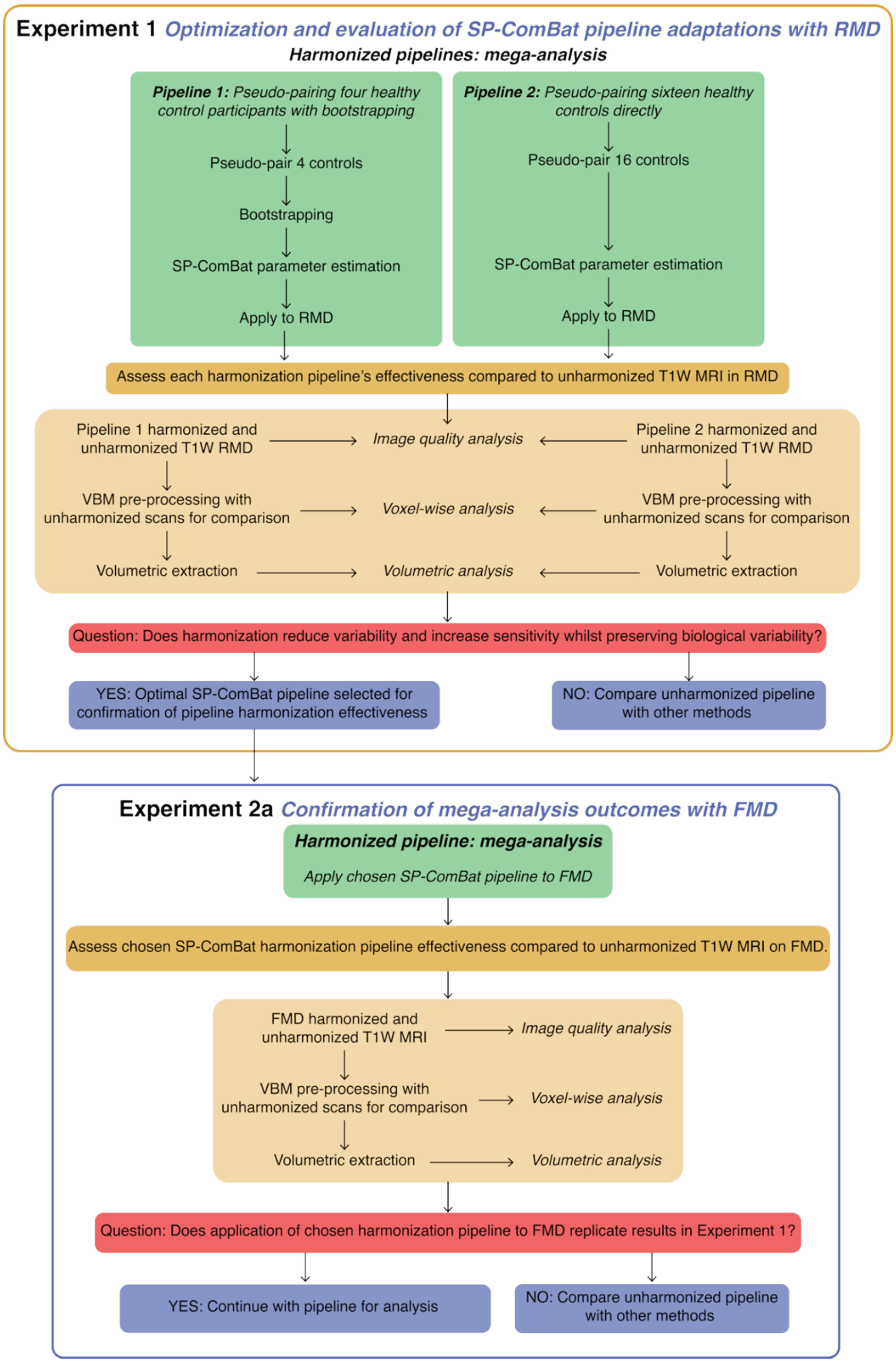


*Supplementary Figure 1: Pre-registered study design for unpaired SP-ComBat harmonization pipelines. The study was pre-registered prior to analysis and reports results from Experiments 1 and 2a. Experiment 1 involved developing and optimizing two SP-ComBat pipelines adapted for cross-sectional unpaired data and evaluating their ability to harmonize multi-study T1W MRI in the absence of traveling subjects. Experiment 2a applied the pipeline selected in Experiment 1 to the full multi-study dataset (FMD) to replicate and further assess harmonization effectiveness. The go/no-go decision point is indicated by the red box. FMD, full multi-study dataset; MRI, magnetic resonance imaging; RMD, representative multi-study dataset; SP-ComBat, Superpixel-ComBat; T1W, T1-weighted; VBM, voxel-based morphometry*

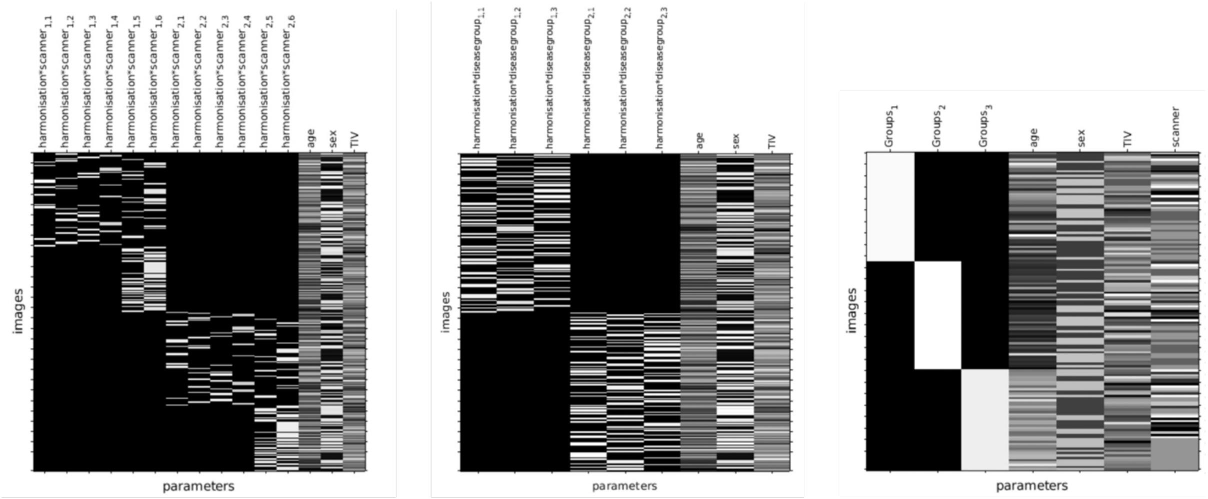


**A** Example harmonization x scanner flexible factorial design matrix

**B** Example harmonization x group flexible factorial design matrix

**C** Example disease group ANOVA design matrix

*Supplementary Figure 2: SPM12 general linear model design matrices for voxel-wise harmonization and disease group analyses. Flexible factorial GLM design matrices for comparing unharmonized and harmonized grey and white matter TPMs across scanner type (A) and disease group (B), and an example one-way ANOVA design matrix for disease group differences (C). In (A), 'harmonization' denotes unharmonized (1) or harmonized (2) status and 'scanner' denotes TRACK-HD London (1), TRACK-HD Leiden (2), TRACK-HD Paris (3), TRACK-HD Vancouver (4), HD-YAS (5), or HD-CSF (6). In (B), 'harmonization' denotes unharmonized (1) or harmonized (2) status and 'disease group' denotes healthy controls (1), pre-CMD (2), and post-CMD (3). In (C), disease groups are healthy controls (1), pre-CMD (2), and post-CMD (3); scanner was included as a covariate in unharmonized analyses only. HD, Huntington's disease; HD-CSF, HD Cerebrospinal Fluid Study; HD-YAS, HD Young Adult Study; GLM, general linear model; HD, Huntington's disease; post-CMD, pwHD after clinical motor diagnosis; pre-CMD, pwHD before clinical motor diagnosis; TPM, tissue probability map.*

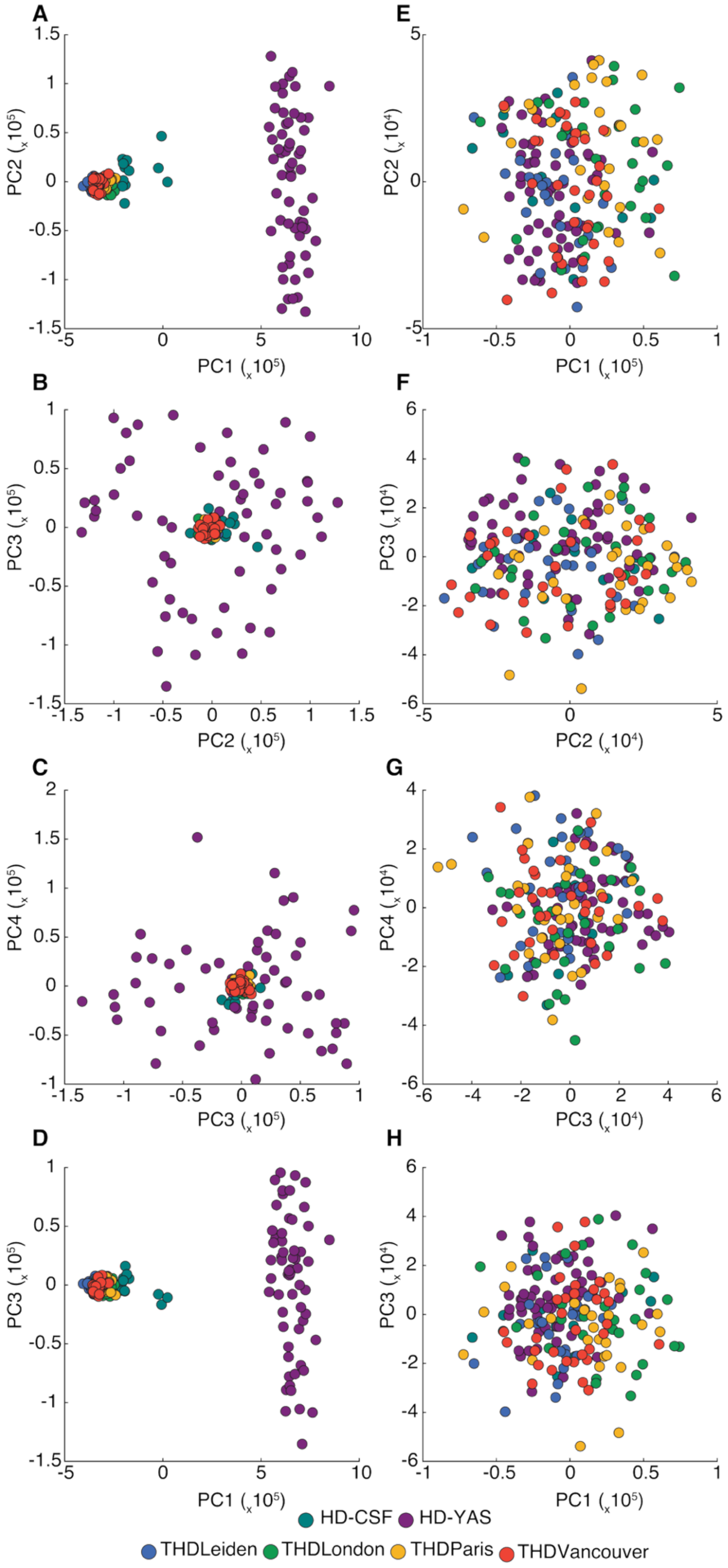


*Supplementary Figure 3: Principal component analysis of intracranially masked voxel intensities in healthy controls before and after SP-ComBat harmonization. PCA applied to intracranially masked voxels from healthy control T1W MRI scans before harmonization (A–D) and after SP-ComBat harmonization (E–H), confirming that scanner-driven clustering observed in the full dataset reflected scanner differences rather than disease-related variance. HD, Huntington's disease; HD-CSF, HD Cerebrospinal Fluid Study; HD-YAS, HD Young Adult Study; PC, principal component; PCA, principal component analysis; THD, TRACK-HD study site.*

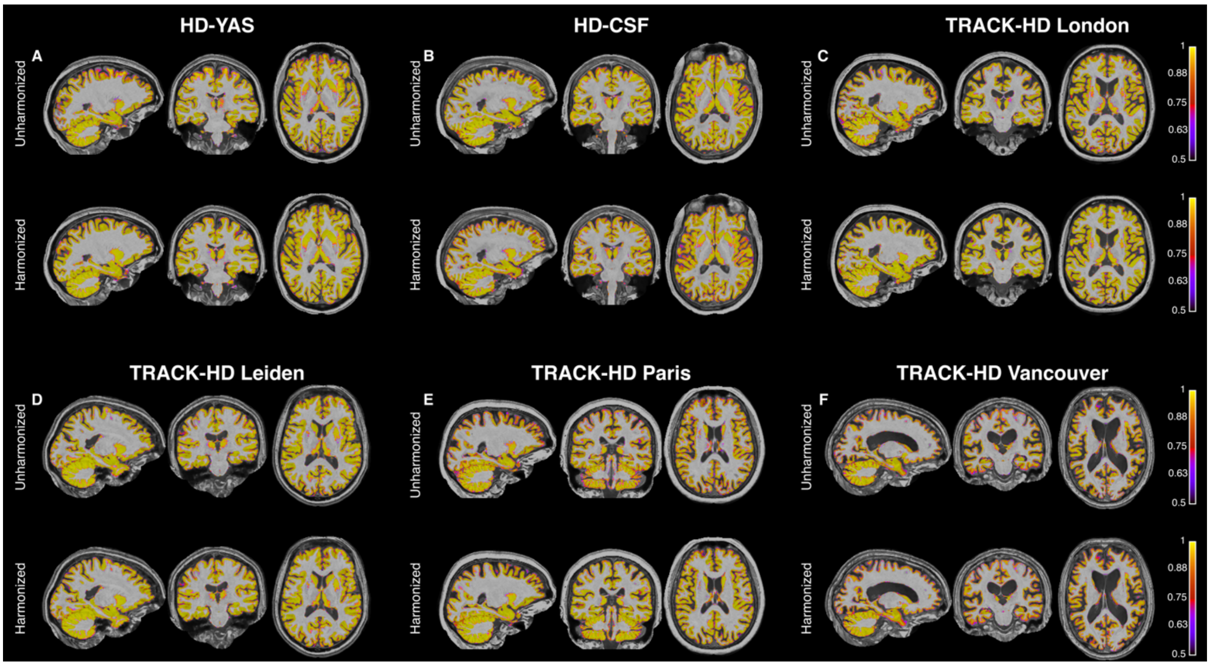


*Supplementary Figure 4: Scanner-specific grey matter segmentations before and after SP-ComBat harmonization across all six scanners. Grey matter segmentations overlaid on native-space T1W MRI for representative scans from HD-YAS (A), HD-CSF (B), TRACK-HD London (C), TRACK-HD Leiden (D), TRACK-HD Paris (E), and TRACK-HD Vancouver (F), shown before and after harmonization. Images are displayed in three-plane orthogonal view with representative sagittal, coronal, and axial slices. Both unharmonized and harmonized segmentations are overlaid on the harmonized T1W MRI to enable direct visual comparison on a common reference image. HD, Huntington's disease; HD-CSF, HD Cerebrospinal Fluid Study; HD-YAS, HD Young Adult Study.*

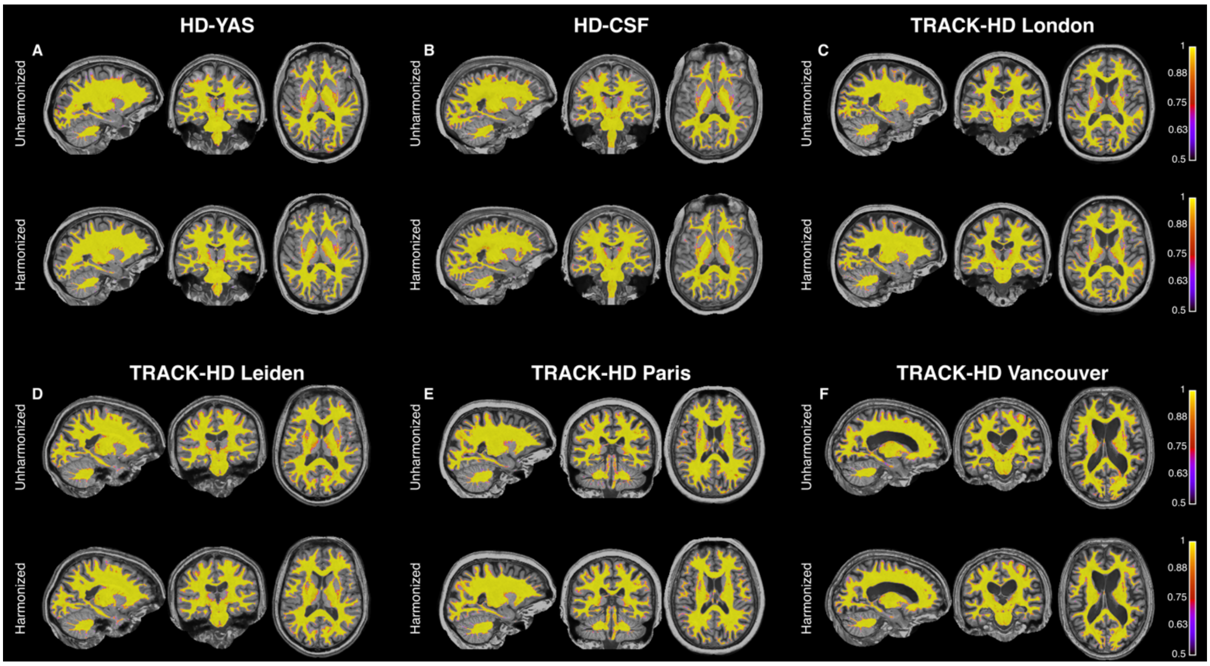


*Supplementary Figure 5: Scanner-specific white matter segmentations before and after SP-ComBat harmonization across all six scanners. White matter segmentations overlaid on native-space T1W MRI for representative scans from HD-YAS (A), HD-CSF (B), TRACK-HD London (C), TRACK-HD Leiden (D), TRACK-HD Paris (E), and TRACK-HD Vancouver (F), shown before and after harmonization. Images are displayed in three-plane orthogonal view with representative sagittal, coronal, and axial slices. Both unharmonized and harmonized segmentations are overlaid on the harmonized T1W MRI to enable direct visual comparison on a common reference image. HD, Huntington's disease; HD-CSF, HD Cerebrospinal Fluid Study; HD-YAS, HD Young Adult Study.*

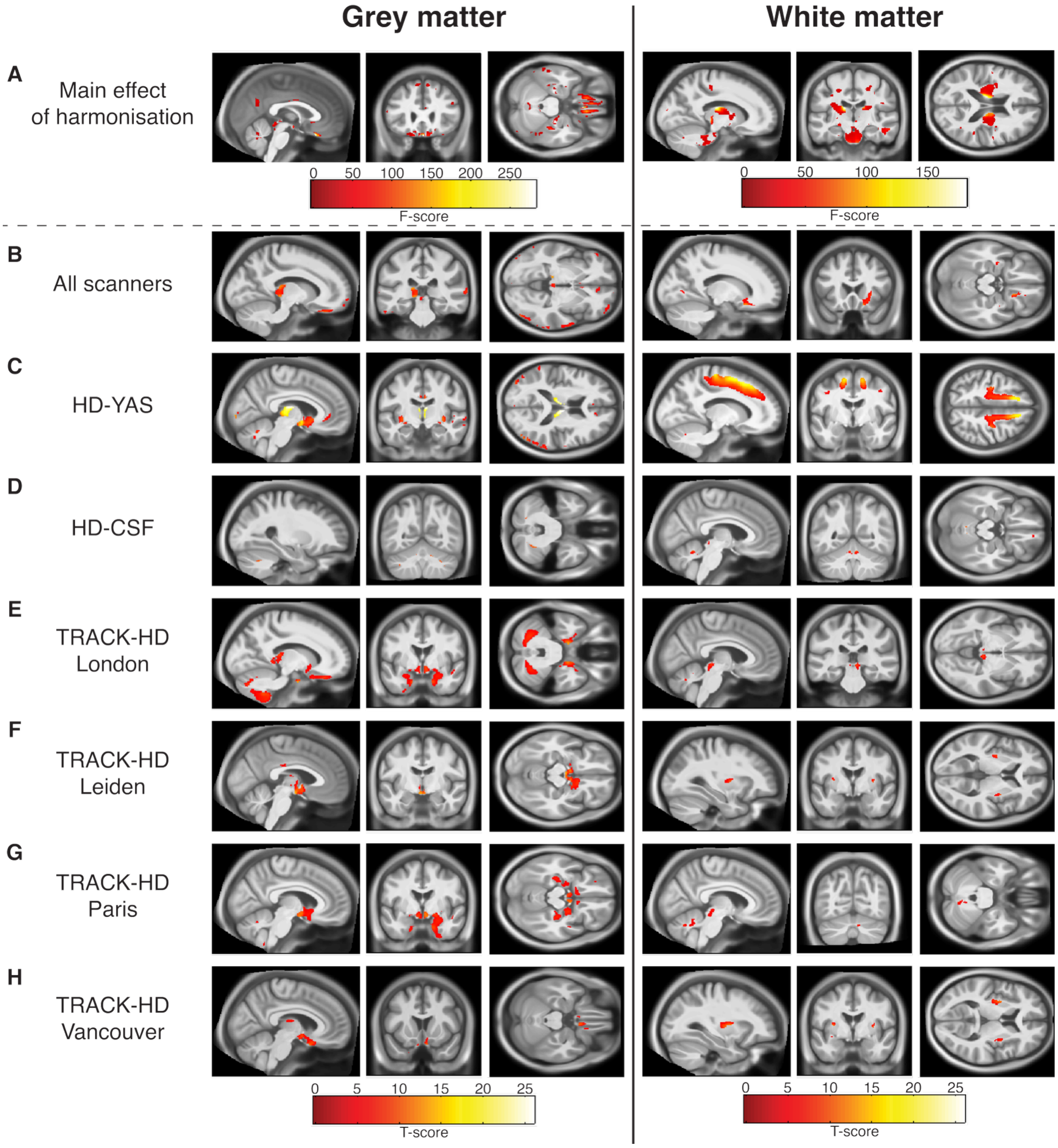


*Supplementary Figure 6: Voxel-wise statistical parametric maps of grey and white matter differences between harmonized and unharmonized TPMs across all scanners. Statistical parametric maps showing grey and white matter differences between harmonized and unharmonized TPMs for the overall F-contrast (A), the overall T-contrast for harmonized > unharmonized (B), and scanner-specific T-contrasts for HD-YAS (C), HD-CSF (D), TRACK-HD London (E), TRACK-HD Leiden (F), TRACK-HD Paris (G), and TRACK-HD Vancouver (H). Significant clusters are overlaid onto sagittal, axial, and coronal sections of the study population-average brain; slices were selected at the contrast global maximum. The main effect of harmonization is displayed as F-scores; scanner-specific contrasts are displayed as T-scores. All comparisons are displayed on the same T-score scale (0–25); threshold p < 0.05, voxel-wise FWE-corrected. FWE, family-wise error; HD, Huntington's disease; HD-CSF, HD Cerebrospinal Fluid Study; HD-YAS, HD Young Adult Study; TPMs, tissue probability maps.*

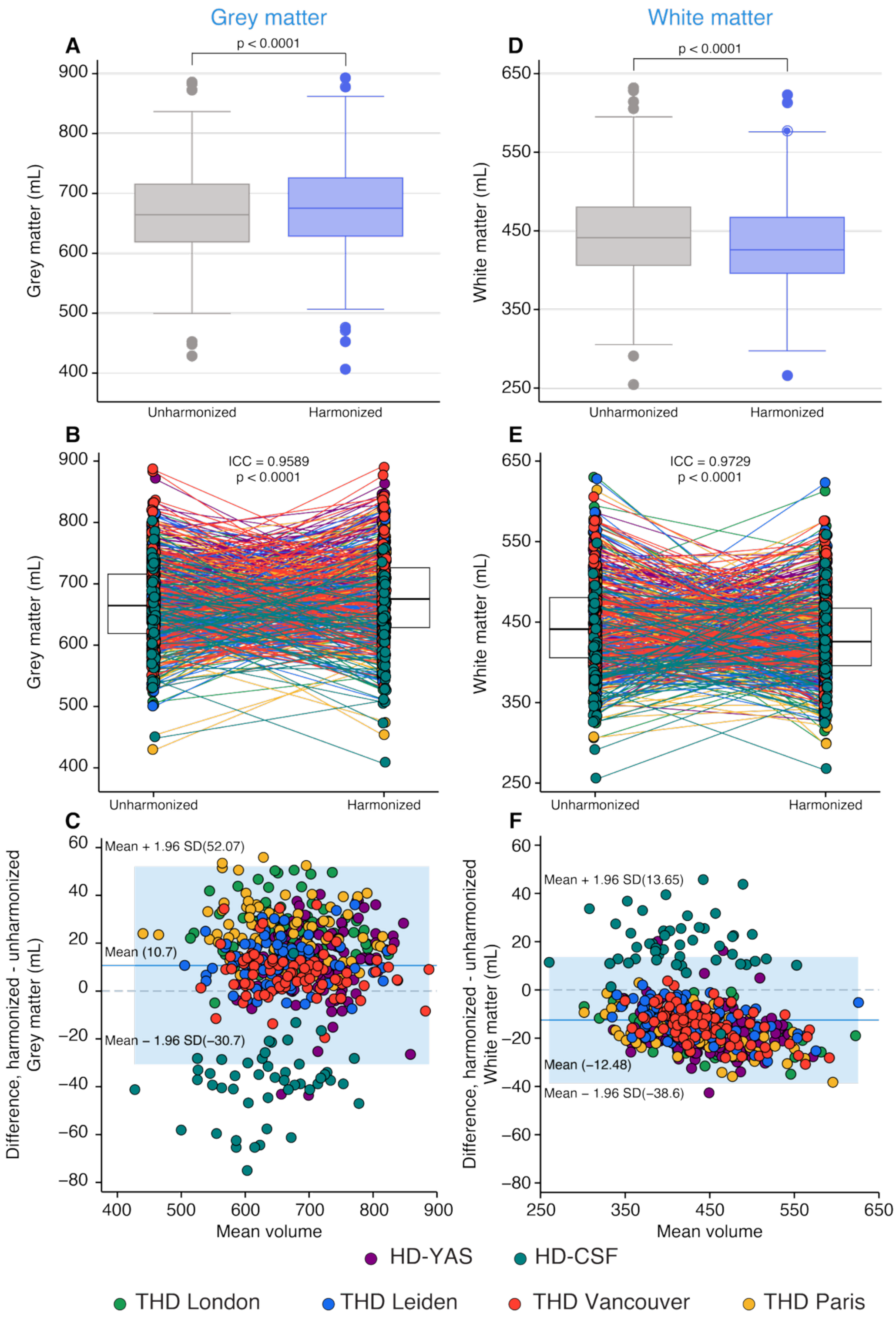


*Supplementary Figure 7: Agreement between unharmonized and harmonized absolute grey and white matter volumes across all scanners. A and D) Differences between unharmonized and harmonized volumes in A) grey and D) whiter matter. P-values obtained from mixed-effects models including harmonization status and its interactions with scanner and disease group as fixed effects, with subject modelled as a random intercept. Boxes show first and third quartiles, the central band shows the median, and the whiskers show data within 1.5 IQR of the median. B and E) Spaghetti plots overlaid on box plots representing the agreement between volumetric measurements before and after harmonization in B) grey, and E) white matter. C and F) Bland-Altman plots for agreement between unharmonized and harmonized volumes in C) grey and F) white matter with the 95% limits of agreement represented by the limits of the light blue shaded region; dark blue lines represent the mean difference; dashed line represents y=0. HD, Huntington's disease; HD-CSF, HD Cerebrospinal Fluid Study; HD-YAS, HD Young Adult Study; ICC, intra-class correlation; THD, TRACK-HD study site; TPM, tissue probability map.*

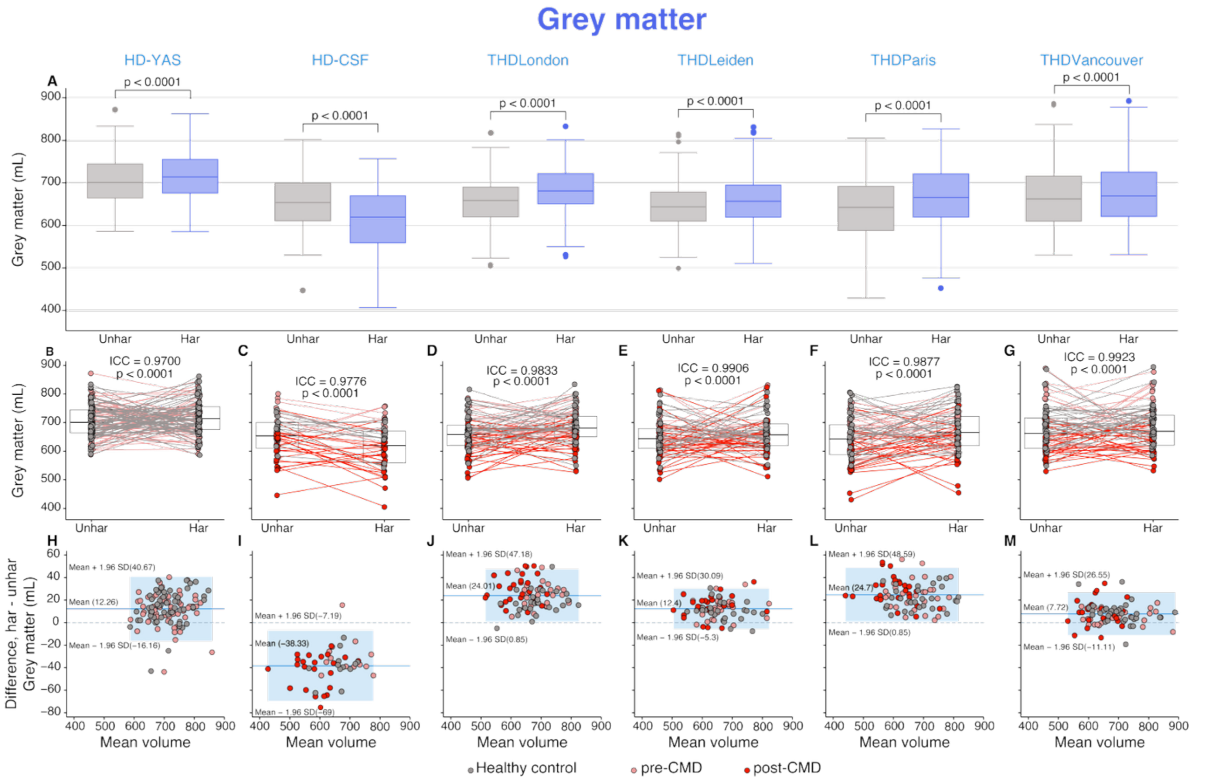


*Supplementary Figure 8: Scanner-specific agreement between unharmonized and harmonized grey matter volumes. A) Differences between unharmonized and harmonized grey matter volumes for each scanner. P-values extracted from Bonferroni-corrected post-hoc pairwise comparisons of estimated marginal means derived from the mixed-effects model (fixed effects: harmonization × scanner × disease group; random intercept: subject). Boxes show first and third quartiles, the central band shows the median, and the whiskers show data within 1.5 IQR of the median. B-G) Spaghetti plots overlaid on box plots representing the agreement between volumetric measurements before and after harmonization in grey matter for B) HD-YAS, C) HD-CSF, D) TRACK-HD London, E) TRACK-HD Leiden, F) TRACK-HD Paris, and G) TRACK-HD Vancouver. H-M) Bland-Altman plots for agreement between unharmonized and harmonized volumes in grey matter for H) HD-YAS, I) HD-CSF, J) TRACK-HD London, K) TRACK-HD Leiden, L) TRACK-HD Paris, and M) TRACK-HD Vancouver with the 95% limits of agreement represented by the limits of the light blue shaded region; dark blue lines represent the mean difference; dashed line represents y=0. Har, harmonized; HD, Huntington's disease; HD-CSF, HD Cerebrospinal Fluid Study; HD-YAS, HD Young Adult Study; ICC, intra-class correlation; post-CMD, pwHD after clinical motor diagnosis; pre-CMD, pwHD before clinical motor diagnosis; THD, TRACK-HD study site; TPM, tissue probability map; unhar, unharmonized.*

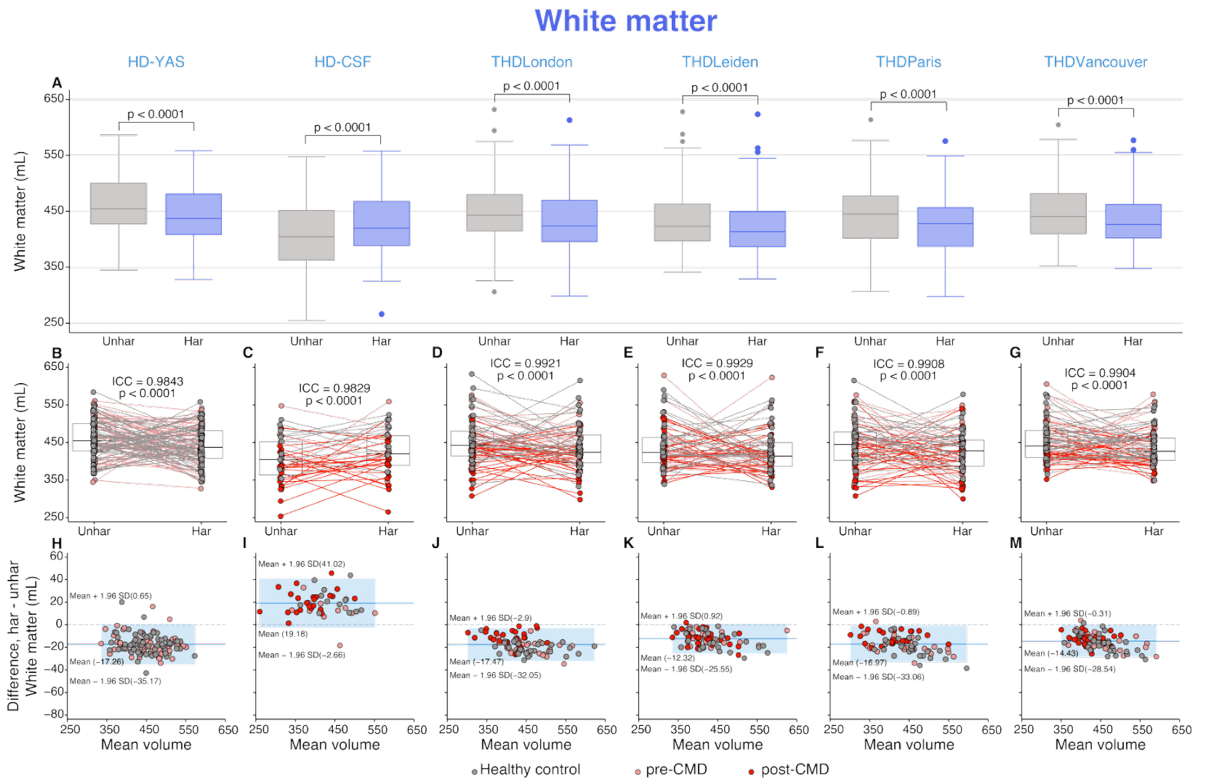


*Supplementary Figure 9: Scanner-specific agreement between unharmonized and harmonized white matter volumes. A) Differences between unharmonized and harmonized grey matter volumes for each scanner. P-values extracted from Bonferroni-corrected post-hoc pairwise comparisons of estimated marginal means derived from the mixed-effects model (fixed effects: harmonization × scanner × disease group; random intercept: subject). Boxes show first and third quartiles, the central band shows the median, and the whiskers show data within 1.5 IQR of the median. B-G) Spaghetti plots overlaid on box plots representing the agreement between volumetric measurements before and after harmonization in grey matter for B) HD-YAS, C) HD-CSF, D) TRACK-HD London, E) TRACK-HD Leiden, F) TRACK-HD Paris, and G) TRACK-HD Vancouver. H-M) Bland-Altman plots for agreement between unharmonized and harmonized volumes in grey matter for H) HD-YAS, I) HD-CSF, J) TRACK-HD London, K) TRACK-HD Leiden, L) TRACK-HD Paris, and M) TRACK-HD Vancouver with the 95% limits of agreement represented by the limits of the light blue shaded region; dark blue lines represent the mean difference; dashed line represents y=0. Har, harmonized; ICC, intra-class correlation; post-CMD, pwHD after clinical motor diagnosis; pre-CMD, pwHD before clinical motor diagnosis; THD, TRACK-HD study site; TPM, tissue probability map; Unhar, unharmonized.*

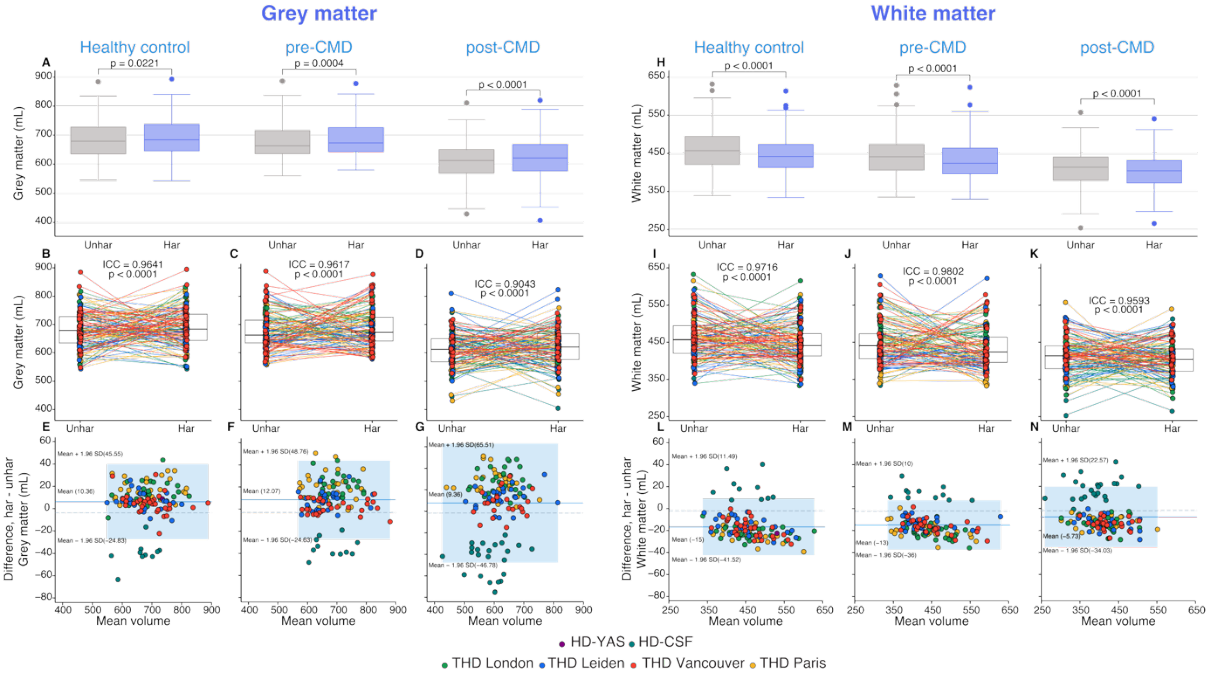


*Supplementary Figure 10: Disease-group-specific agreement between unharmonized and harmonized grey and white matter volumes with QC > 3. A,H) Differences between unharmonized and harmonized grey matter A) and white matter H) volumes for each disease group. P-values extracted from Bonferroni-corrected post-hoc pairwise comparisons of estimated marginal means derived from the mixed-effects model (fixed effects: harmonization × scanner × disease group; random intercept: subject). Boxes show first and third quartiles, the central band shows the median, and the whiskers show data within 1.5 IQR of the median. B-D, I-K) Spaghetti plots overlaid on box plots representing the agreement between volumetric measurements before and after harmonization in grey and white matter for B,I) healthy controls, C,J) pre-CMD, and D,K) post-CMD. E-G,L-N) Bland-Altman plots for agreement between unharmonized and harmonized volumes in grey and white matter for E,L) healthy controls, F,M) pre-CMD, and G,N) post-CMD, with the 95% limits of agreement represented by the limits of the light blue shaded region; dark blue lines represent the mean difference; dashed line represents y=0. Har, harmonized; post-CMD, pwHD after clinical motor diagnosis; pre-CMD, pwHD before clinical motor diagnosis; TPMs, tissue probability maps; Unhar, unharmonized.*
